# Temperature sensitivity of the interspecific interaction strength of coastal marine fish communities

**DOI:** 10.1101/2022.06.02.494625

**Authors:** Masayuki Ushio, Tetsuya Sado, Takehiko Fukuchi, Sachia Sasano, Reiji Masuda, Yutaka Osada, Masaki Miya

**Author notes:** Email: MU,; TS,; TF,; SS,; RM,; YO,; MM. **Data availability statement:** Source scripts for the analyses and figure generations are archived in Zenodo (https://doi.org/10.5281/zenodo.7865958) and publicly available at Github (https://github.com/ong8181/eDNA-BosoFish-network). DNA data is deposited DDBJ Sequence Read Archive (DRA submission ID = DRA014111).

## Abstract

The effects of temperature on interaction strengths are important for understanding and forecasting how global climate change impacts marine ecosystems; however, tracking and quantifying interactions of marine fish species is practically difficult especially under field conditions, and thus, how temperature influences their interaction strengths under field conditions remains poorly understood. We herein performed quantitative fish environmental DNA (eDNA) metabarcoding on 550 seawater samples that were collected twice a month from 11 coastal sites for two years in the Boso Peninsula, Japan, and analyzed eDNA monitoring data using nonlinear time-series analytical tools. We detected fish-fish interactions as information flow between eDNA time series, reconstructed interaction networks for the top 50 frequently detected species, and quantified pairwise, fluctuating interaction strengths. Although there was a large variation, water temperature influenced fish-fish interaction strengths. The impact of water temperature on interspecific interaction strengths varied among fish species, suggesting that fish species identity influences the temperature effects on interactions. For example, interaction strengths that *Halichoeres tenuispinis* and *Microcanthus stringatus* received strongly increased with water temperature, while those of *Engraulis japonicus* and *Girella punctata* decreased with water temperature. An increase in water temperature induced by global climate change may change fish interactions in a complex way, which consequently influences marine community dynamics and stability. Our research demonstrates a practical research framework to study the effects of environmental variables on interaction strengths of marine communities in nature, which would contribute to understanding and predicting natural marine ecosystem dynamics.

## Introduction

Interspecific interactions are key to understanding and predicting the dynamics of ecological communities (May, 1972; Tang et al., 2014; Wootton and Emmerson, 2005; Wootton and Stouffer, 2015). Theoretical and empirical studies have shown that various properties of interspecific interactions, namely, the number of interactions, sign (positive or negative), strength, and correlations of pair-wise interactions (e.g., predator-prey interactions), influence the dynamics, stability, and diversity of ecological communities in terrestrial and aquatic ecosystems (Allesina et al., 2015; Mougi and Kondoh, 2012; Ratzke et al., 2020; Tang et al., 2014; Ushio, 2022a; Ushio et al., 2018a). Interaction strength is a fundamental property, and previous studies examined the relationship between interaction strengths and ecological community properties (Wootton and Emmerson, 2005; Wootton and Stouffer, 2015). The dominance of weak interactions stabilizes the dynamics of a natural fish community (Ushio et al., 2018a). Interaction strengths have been reported to decrease with increases in the diversity of experimental microbial communities (Ratzke et al., 2020), and this holds true even for more diverse ecological communities under field conditions (Ushio, 2022a).

Environmental variables exert significant effects on interspecific interaction strengths. The effects of temperature on interaction strengths are important for understanding and forecasting the impact of the ongoing global climate change on ecosystems. The relationship between temperature and interaction strengths has been investigated in terrestrial and aquatic ecosystems for decades (Adams and Zhang, 2009; Allan et al., 2015; Coley and Aide, 1991; Hein et al., 2014; Kishi et al., 2005; Kordas et al., 2011; Kratina et al., 2012; Rall et al., 2010; Thakur et al., 2017). In terrestrial ecosystems, Coley & Aide (1991) proposed that interactions between plants and insect herbivores were generally stronger in warmer regions, which may result in stronger negative density dependence for tree species and contribute to higher plant diversity in a tropical region (Forrister et al., 2019). The influence of temperature on interaction strengths has also been investigated in other systems: terrestrial arthropods interactions (Rall et al., 2010), fish-fish interactions (Allan et al., 2015; Hein et al., 2014), and fish-prey interactions (Kishi et al., 2005). In addition, Wieczynski *et al*. (2021) showed that species-level functional traits (e.g., body size and shape) may underlie the relationships between temperature and interaction strengths. Therefore, interplays among temperature, interaction strengths, species identity, and community-level function (e.g., primary production) is prevalent, and a more detailed understanding of the complex interactions among them is critical for predicting ecosystem responses to climate change.

In marine ecosystems, interspecific interactions in ecological communities play a fundamental role in system dynamics, such as population dynamics, primary production, and nutrient cycling (Hannisdal et al., 2017; Penn et al., 2019; Ushio et al., 2018a), as well as ecosystem services, including food supply (Smith et al., 2019). Although many studies have evaluated the effects of temperature on interaction strengths of marine organisms (Kordas et al., 2011), most of these studies have been performed targeting relatively immobile or small organisms (Bertness and Ewanchuk, 2002; Chen et al., 2012) or under laboratory conditions (e.g., mesocosm experiments; Allan et al., 2015). While the previous studies have provided invaluable information, it currently remains unclear how temperature influences the strengths of interspecific interactions of larger, more mobile marine organisms, such as fish, which may exert strong top-down regulations on ecological communities, especially under field condition. This may be due to the difficulties associated with detecting and measuring interaction strengths among multiple, relatively large, mobile species under field conditions; the identification of quickly-moving fish species and the quantification of their abundance under field conditions are challenging, and the quantification of their interactions is even more difficult. Overcoming these difficulties and understanding how temperature influences the strength of interspecific interactions of marine fish communities will provide insights into how marine fish communities are assembled, how they may respond to the ongoing and future global climate change, and how the changes in fish communities may transmit to other trophic levels.

Environmental DNA (eDNA), defined here as extra-organismal DNA left behind by macro-organisms (Bohmann et al., 2014), has been attracting increasing attention as an indirect genetic marker for inferring the presence of species for biodiversity monitoring (Cristescu and Hebert, 2018; Deiner et al., 2017). A simple protocol for collecting eDNA samples from aquatic environments facilitates continuous biodiversity monitoring at multiple sites (Deiner et al., 2017), and eDNA metabarcoding (the simultaneous detection of multiple species using universal primers and a high-throughput sequencer) provides useful information on the dynamics of ecological communities (Bálint et al., 2018; Miya, 2022; Miya et al., 2020; Ushio, 2022a). Recent studies demonstrated that eDNA metabarcoding combined with frequent water sampling enabled the efficient monitoring of high-diversity ecological communities (Bista et al., 2017; Djurhuus et al., 2020; Ushio, 2022a). Furthermore, if eDNA metabarcoding is performed quantitatively (e.g., by including spike-in DNAs; Ushio et al., 2018b), these data may contain information on species abundance.

More importantly, recent studies have shown that information on interspecific interactions may be embedded in multispecies eDNA time series, particularly when the data is “quantitative” (Ushio, 2022a). Advances in nonlinear time series analyses have enabled the quantification of interspecific interactions only from time series data. For example, transfer entropy (TE) is a method based on the information theory that quantifies information flow between two variables (Runge et al., 2012; Schreiber, 2000). Information flow can be an index of interspecific interactions when applied to ecological data. Convergent cross mapping (CCM) is a causality detection tool in empirical dynamic modeling (Sugihara et al., 2012), which is based on the dynamical theory. TE and CCM have more recently been understood under the information theory framework, and unified information-theoretic causality (UIC) may also quantify information flow between variables (Osada et al., 2023). In addition, an improved version of a sequentially-weighted global linear map (S-map), called the multiview distance regularized (MDR) S-map, enables accurate quantifications of interaction strengths of a large interaction network even when the number of network nodes exceeds the time series length (Chang et al., 2021). These advanced statistical tools facilitate the detection and quantification of interspecific interactions from quantitative, multispecies eDNA time-series data.

In the present study, we collected eDNA samples twice a month from 11 coastal sites at the southern tip of the Boso Peninsula, central Japan (Fig. 1a) for two years (a total of 550 samples). This region is located on the Pacific side of Honshu Island around 35°N and is markedly affected by the warm Kuroshio Current, cold Oyashio Current, and inner-bay water from Tokyo Bay. These geographic and oceanographic characteristics form latitudinal temperature gradients, allowing us to investigate temperature effects on interspecific interactions. We obtained quantitative eDNA metabarcoding data by combining MiFish eDNA metabarcoding data (relative abundance data) and the eDNA concentration data obtained through quantitative PCR (qPCR) for one of the most common fish species, *Acanthopagrus schlegelii* (e.g., absolute abundance data of *A. schlegelii* eDNA was utilized as an internal spike-in DNA; see Methods). Based on quantitative, multispecies eDNA time-series data, pairwise species interactions were detected as information flow between species using a nonlinear time series analysis (Osada et al., 2023). In addition, interaction strengths at each time point were quantified for the detected fish-fish interactions using the MDR S-map method (Chang et al., 2021). As our eDNA time series was taken twice a month, the interactions detected should also have the same time scale (e.g., the interactions detected may cause changes in the population size at the same time scale), which means that we tend to focus on behavior-level interactions (e.g., feeding, vigilance, and schooling) rather than birth-death process in the present study (except for predation). In addition, the interactions we detected by the time series analysis could include various types of interactions such as competition and mutualism that could be involved in the population dynamics. We hypothesized that (1) a positive relationship exists between temperature and interaction strengths at the community level, as found in other ecosystems and many types of interactions, and (2) the relationship between the patterns of temperature and interaction strengths varies among fish species because of species-specific ecologies and the behavior of component species.

**Figure 1.**
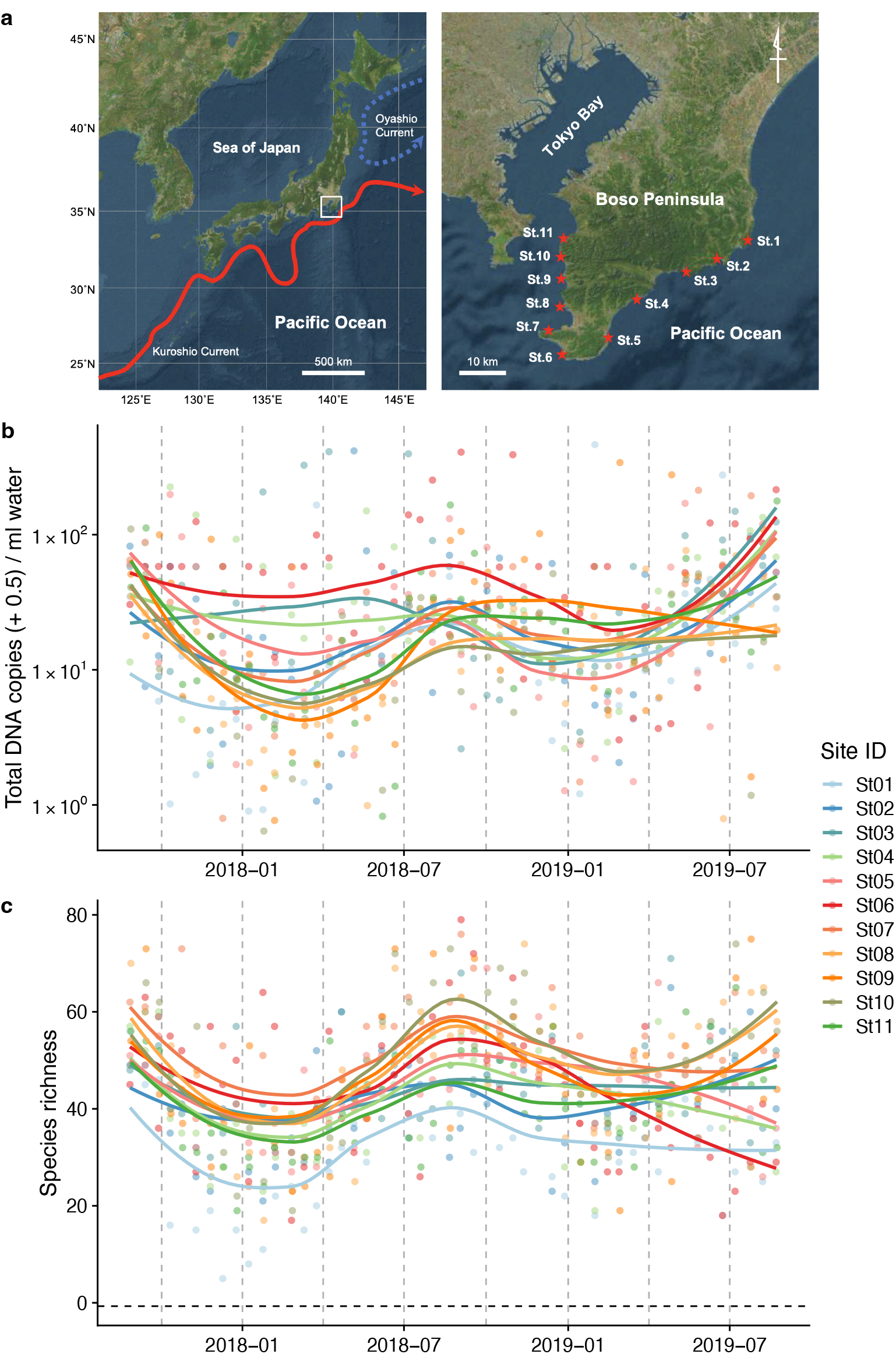
Study sites and overall dynamics of environmental DNA (eDNA) concentrations and the number of fish species detected. (**a**) Study sites in the Boso Peninsula. The study sites are influenced by the Kuroshio current (red arrow; left panel) and distributed along the coastal line in the Boso Peninsula (right panel). (**b**) Total eDNA copy numbers estimated by quantitative eDNA metabarcoding (see Methods for detail). (**c**) Fish species richness detected by eDNA metabarcoding. Points and lines indicate raw values and LOESS smoothing lines, respectively. The line color indicates the sampling site. Warmer colors generally correspond to study sites with a higher mean water temperature.

## Materials and Methods

### Study sites and environmental observations

We selected 11 sites (Stations [Sts.] 1–11) for seawater sampling along the shoreline of the southern tip of the Boso Peninsula (Fig. 1a). The 11 stations are located within or surrounded by rocky shores with intricate coastlines. They are arranged such that the five stations on the Pacific and Tokyo Bay sides are approximately at the same latitudinal intervals, with St. 6 in between. The northernmost stations on both sides (Sts. 1 and 11) are markedly affected by the cold Oyashio Current (Yang et al., 1993) and inner-bay water from Tokyo Bay (Fujiwara and Yamada, 2002), respectively, while the remaining stations (Sts. 2–10) are markedly affected by the warm Kuroshio Current flowing northward along the Pacific coast as well as its branches (Soh, 2003). Before seawater sampling at each station, we measured seawater temperature (°C) and salinity (‰) using a portable water-quality meter (WQC-30, Toa DKK, Tokyo, Japan). In addition, we recorded the sampling start time, latitude/longitude, weather and sea conditions, and turbidity.

### Collection of eDNA samples

Detailed information on seawater sampling and on-site filtration is provided in Supporting Information. We collected seawater samples twice a month (i.e., around the first and second quarter moons) for two years between August 2017 and August 2019 by casting a bucket fastened to a rope at the 11 sites. We filtered the collected seawater using a combination of a filter cartridge and disposable syringe until the final filtration volume reached 1,000 mL for each replicate and obtained duplicate samples at each site. We added RNAlater (Thermo Fisher Scientific, DE, USA) into the cartridge to prevent eDNA degradation. We made a filtration blank (FB) by filtering 500 mL of purified water in the same manner at the end of each day of water sampling.

### Laboratory protocol

Detailed information on the laboratory protocol is provided in Supporting Information. We thoroughly sterilized the workspace and equipment in the DNA extraction, pre-PCR, and post-PCR rooms before all experiments. We used filtered pipette tips and conducted all eDNA extraction, pre-PCR, and post-PCR manipulations in three different dedicated rooms that were separate from each other to safeguard against cross-contamination.

We extracted eDNA from filter cartridges using the DNeasy Blood & Tissue kit (Qiagen, Hilden, Germany) following the methods developed by Miya *et al.* (2016) with slight modifications (Minamoto et al., 2021; Miya and Sado, 2019a). After removing RNAlater from filter cartridges, they were subjected to lysis using proteinase K. We purified the collected DNA extracts using the above kit following the manufacturer’s protocol and the final elution volume was set to 200 µL. An extraction blank (EB) was also created during this process.

We employed two-step PCR for paired-end library preparation using the MiSeq platform (Illumina, CA, USA). We generally followed the methods developed by Miya et al. (2015) and subsequently modified by Miya & Sado (2019b). We performed the first-round PCR (1st PCR) using a mixture of the following six primers: MiFish-U-forward, MiFish-U-reverse, MiFish-E-forward-v2, MiFish-E-reverse-v2, MiFish-U2-forward, and MiFish-U2-reverse (Table 1). The 1st PCR amplified a hypervariable region of the mitochondrial 12S rRNA gene (*ca*. 172 bp; hereafter called the “MiFish sequence”). We also prepared a 1st PCR blank (1B) during this process, in addition to FB and EB. After the 1st PCR, PCR products were purified, quantified, diluted to 0.1 ng/µL, and used as templates for the second-round PCR (2nd PCR).

**Table 1.**
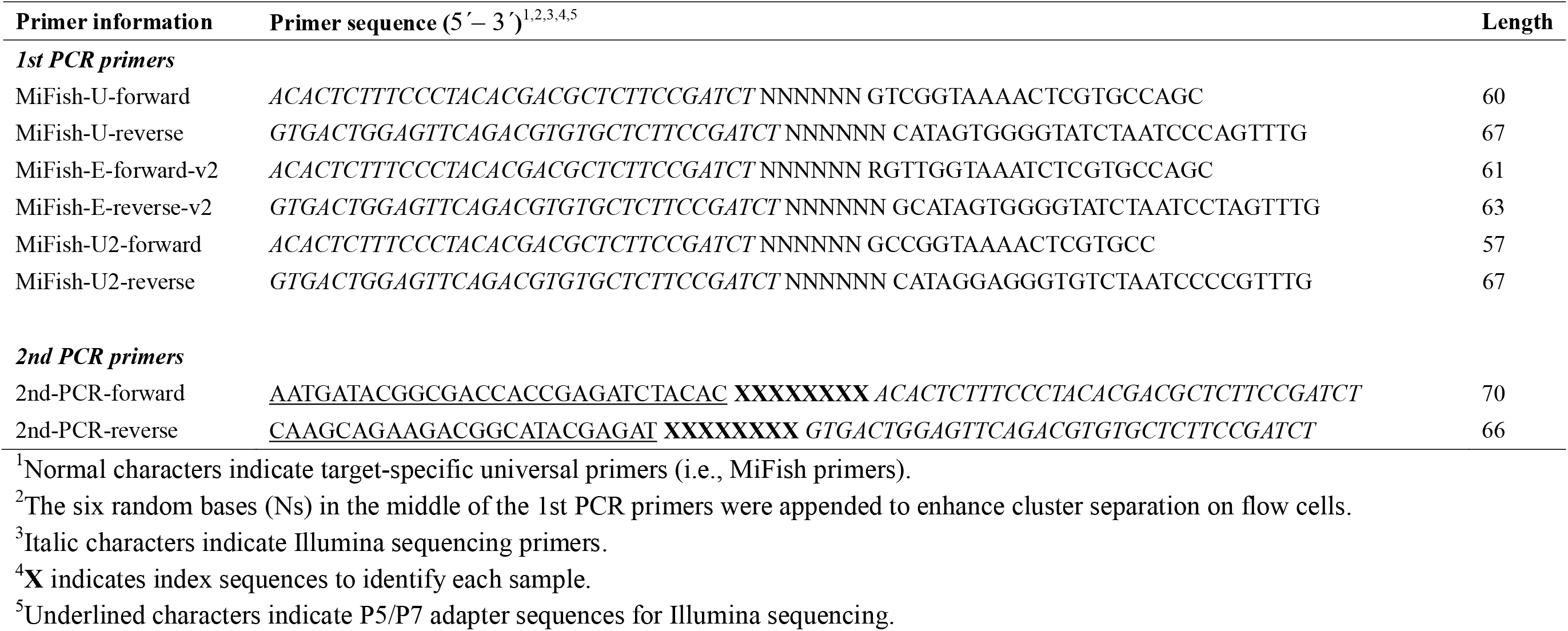
Primer sequences used in the present study.

We performed the 2nd PCR to append dual-indexed sequences and flow cell-binding sites for the MiSeq platform. We prepared a 2nd PCR blank (2B) during this process in addition to FB, EB, and 1B. In total, we made 195 blanks (FB = 60, EB = 50, 1B = 50, 2B = 35) to monitor contamination during the on-site filtration, subsequent DNA extraction, and 1st and 2nd PCR of 550 samples.

We pooled each of the individual paired-end libraries in an equal volume, performed electrophoresis on an agarose gel (Invitrogen, CA, USA), and excised the target amplicons (*ca*. 370 bp). The concentrations of the size-selected libraries were measured, diluted to 10–12.0 pM, and sequenced on the MiSeq platform using the MiSeq v2 Reagent Kit for 2 × 150 bp PE (Illumina, CA, USA) following the manufacturer’s protocol. We performed 21 MiSeq runs (with eDNA samples of other projects) to complete the analysis, which generated 71,892,685 reads in total for the eDNA samples of this project.

### Sequence analysis

We performed data preprocessing and analyses of raw MiSeq reads from the MiSeq run using PMiFish ver. 2.4 (https://github.com/rogotoh/PMiFish; Miya et al. 2020). Detailed information on sequence data processing is provided in Supporting Information. Forward (R1) and reverse (R2) reads were merged while discarding short reads (<100 bp) after tail trimming and paired reads with too many differences (>5 positions) in the aligned region (*ca.* 65 bp). Primer sequences were removed from merged reads and reads without subjecting the primer sequences to quality filtering to remove low quality reads. Preprocessed reads were dereplicated, and all singletons, doubletons, and tripletons were removed from subsequent analyses to avoid false positives (Edgar, 2010). Dereplicated reads were denoised to generate amplicon sequence variants (ASVs) that removed all putatively chimeric and erroneous sequences (Callahan et al., 2017; Edgar, 2016).

ASVs were subjected to taxon assignments to species names with a sequence identity of >98.5% with the reference sequences (two nucleotide differences allowed) and a query coverage of ≥90%. ASVs with identical fish species names were clustered as molecular operational taxonomic units (MOTUs) following the procedure described in Supporting Information. MiFish DB ver. 43 was used for taxon assignment, comprising 7,973 species distributed across 464 families and 2,675 genera. The ASV table was rarefied to the approximate minimum read number (*ca*. 20,000 reads), which resulted in 10,984,789 reads in total for the rarefied ASV table. At present, the reference sequences are available for about 70% of 4,500 fish species in Japan. However, due to the unknown degree of intraspecific variation, using a uniform threshold of 98.5% to delineate species can result in over-splitting or over-clustering MOTUs. To solve this issue, manual refinement of the taxon assignments was performed based on the phylogenetic tree. To refine the above taxon assignments, family-level phylogenies were reproduced from MiFish sequences from MOTUs and reference sequences (contained in MiFish DB ver. 43) belonging to these families. A total of 103 family-level trees were visually inspected and taxon assignments were revised. The final list of detected species is provided in Table S1. The negative controls produced negligible reads and all of the reads were assigned to non-target taxa. Therefore, we discarded the sequence reads from the negative control samples (see Results and Discussion for details).

### Estimation of DNA copy numbers

The nonlinear time series analytical tools used in the present study require quantitative time-series data. To estimate fish eDNA concentrations, we initially measured the eDNA concentrations of the most common fish species in the region, Japanese Black Seabream (*A. schlegelii*), using qPCR, which was detected in 504 out of 550 seawater samples (91.6%). By dividing *A. schlegelii* sequence reads by the *A. schlegelii* eDNA quantity in each sample, we estimated the number of sequence reads generated per *A. schlegelii* eDNA copy for each sample, and we applied the same conversion factor (i.e., sequence reads /*A. schlegelii* eDNA copy) to other fish species. Note that sequence reads generated per eDNA copy may vary depending on factors such as the level of PCR inhibition and PCR amplification efficiency; however, in general, this “internal spike-in DNA” method has been shown to reasonably estimate the quantity of eDNA concentrations (Ushio, 2022a; Ushio et al., 2018b). More importantly, species-specific biases that may be introduced in the internal spike-in DNA method do not cause serious biases in the outcomes of our nonlinear time-series analysis because it standardizes time series to have a zero mean and a unit variance before analyses and also only utilizes the fluctuation patterns of time series. When we did not detect any *A. schlegelii* eDNA by qPCR, we replaced the “zero” value with the minimum eDNA copy numbers of *A. schlegelii* (0.346 copies/µl DNA). Similarly, when we did not detect any *A. schlegelii* eDNA sequence reads by metabarcoding, we replaced the “zero” value with the minimum eDNA sequence reads of *A. schlegelii* (12 reads/sample). These corrections estimated how many sequence reads were generated per eDNA copy for all samples. Details are provided in Supporting Information.

### Nonlinear time series analysis to detect fish-fish interactions

We detected fish-fish interactions using a nonlinear time series analysis based on quantitative eDNA time-series data from multiple species and sites. Since the reliable detection of interspecific interactions requires sufficient information in the target time series, we selected the top 50 most frequently detected species and excluded other rarer fish species from the analysis. We quantified information flow between two fish species’ eDNA time series by the “unified information-theoretic causality (UIC)” method (Osada et al., 2023) implemented in the “rUIC” v0.1.5 package (Osada and Ushio, 2021) of R (R Core Team, 2022). UIC tests the statistical clarity of information flow between variables in terms of TE (Schreiber, 2000) computed by nearest neighbor regression based on time-delay embedding of explanatory variables (i.e., cross mapping; Sugihara *et al.* 2012). In contrast to the standard method used to measure TE, UIC quantifies information flow as follows:

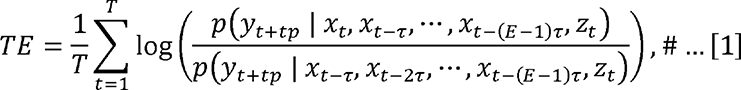

where *x*, *y*, and *z* represent a potential causal variable, effect variable, conditional variable (if available), respectively. *p*(*A*|*B*) represents conditional probability: the probability of *A* conditioned on *B*. *t*, *tp*, τ, and *E* represent the time index, time step, a unit of time lag, and the optimal embedding dimension, respectively. *T* is the total number of points in the reconstructed state space (this is equivalent to the total number of time points – the optimal embedding dimension + 1). For example, if *tp* = –1 in Eqn. [1], UIC tests the causal effect from *y_t_*_–1_ to *x_t_*. Optimal *E* was selected by measuring TE as follows:

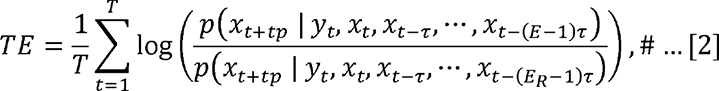

where *E_R_* (< *E*) is the optimal embedding dimension of lower dimensional models. In the present study, the causality time-lag (*tp*) up to –6 (equivalent to three-month time-lag) was tested. Significance tests were conducted by bootstrapping data after embedding (the significance levels were set to 0.05). Eqn. [2] is a TE version of simplex projection (Sugihara and May, 1990). TE measured according to Eqn. [1] gains the advantage of previous causality tests, *i.e.*, standard TE methods (Runge et al., 2012; Schreiber, 2000) and convergent cross mapping (CCM) (Sugihara et al., 2012), the algorithm of which is explained in Osada et al. (2023) and implemented in Osada & Ushio (2021).

By using UIC, we quantified TE between fish eDNA time series. As environmental variables, water temperature (°C), salinity (‰), wave height (m), and tide level (cm) were considered as conditional variables. If the environmental variables had statistically clear influence on fish eDNA dynamics, they were included in the calculation of TE as *z_t_* in Eqn. [1]. This means that the effects of the environmental variables on the fish eDNA abundance were removed in the analysis when detecting interspecific interactions. Importantly, in most cases, water temperature had statistically clear influence on fish eDNA dynamics and included as a conditional variable (*z_t_*) in the embedding. In addition, although water temperature showed clear seasonality in the region (Fig. S1), including water temperature as a conditional variable (*z_t_*) took the effect of the seasonality in detecting causation into account. We merged all eDNA time series across the 11 study sites and standardized the eDNA time series to have zero means and a unit of variance before the analysis. If TE between two fish eDNA time series was statistically clearly higher than zero, we interpreted it as a sign of an interspecific interaction. After the detections of interspecific interactions among the top 50 fish species, we visualized the interaction network (i.e., the reconstruction of fish-fish interaction network). Importantly, UIC quantifies the average information flow between two time series (see Eqn. [1]), and thus there is only one TE value for each pair of fish species, which is critically different from interaction strengths quantified by the MDR S-map (see the following section).

### Nonlinear time series analysis to quantify interaction strengths

We quantified fish-fish interaction strengths for the interspecific interactions detected by UIC. For this purpose, we used an improved version of the sequential locally-weighted global linear map (S-map) (Sugihara, 1994), called the multiview distance regularized S-map (MDR S-map) (Chang et al., 2021). Consider a system that has *E* different interacting variables, and assume that the state space at time *t* is given by *x*(*t*) = {*x*_1_(*t*), *x*_2_(*t*), …, *x_E_*(*t*)}. For each target time point *t*\*, the S-map method produces a local linear model that predicts the future value *x*_1_(*t*\*+*p*) from the multivariate reconstructed state space vector *x*(*t*\*). That is,

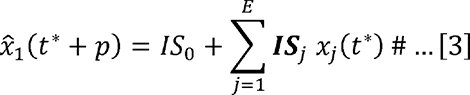

Where 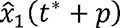 is a predicted value of *x*_1_ at time *t*\*+*p*, *IS_j_* is a regression coefficient and interpreted as interaction strength (or called S-map coefficient), and *IS*_0_ is an intercept of the linear model. The linear model is fit to the other vectors in the state space. However, points that are close to the target point, *x*(*t*\*), are given greater weighting (i.e., locally weighted linear regression). In the standard S-map, the distances between the target point, *x*(*t*\*), and other points are measured by Euclidean distance. However, Euclidean distance cannot be a good measure if the dimension of the state space is high, and in this case, it is impossible to identify nearest neighbors correctly. In the MDR S-map, the distance between the target point 11 and other points are measured by the multiview distance (Chang et al., 2021), which can be calculated by ensembling distances measured in various low-dimensional state spaces (multiview embedding; see Ye and Sugihara, 2016). In addition, to reduce the possibility of overestimation and to improve forecasting skill, regularization (i.e., ridge regression) is also applied (Cenci et al., 2019). Chang et al. (2021) showed that the MDR S-map outperformed other S-map methods and it enables improved estimations of interaction strengths. Importantly, the MDR S-map quantifies interaction strength at each time point (i.e., the maximum possible number of interaction strength values for each fish-fish interaction is the number of time points – the optimal embedding dimension + 1). As in the UIC analysis, we included temperature or other environmental variables as conditional variables if they had statistically clear influence on eDNA dynamics of a particular fish species. In the present study, we implemented the MDR S-map in our custom R package, “macam” v0.0.10 (Ushio, 2022b) and used it for computation.

### Quantification of the temperature sensitivity of fish species interactions

After reconstructing fish interaction networks and quantifying interaction strengths, we analyzed the relationships among fish species interaction strengths and biotic and abiotic variables. We performed all analyses using R v4.2.1 (R Core Team, 2022). Generalized additive mixed models (GAMMs) were performed using the “mgcv” package of R (Wood, 2004). We visualized results with “ggplot2” (Wickham, 2009) and “cowplot” (Wilke, 2017). In the present study, we used the term “statistical clarity” instead of “statistical significance” to avoid misinterpretations, according to the recommendations by Dushoff et al. (2019).

In the first analysis, the relationships between fish-fish interaction strengths and the characteristics of the study sites were analyzed by GAMM. In the analysis, in-strength (interaction strengths that a species receives) and out-strength (interaction strengths that a species gives) were separately calculated. In GAMM, in-strength or out-strength were an explained variable, and an environmental or ecological variable was an explaining variable. Explaining variables included water temperature, species richness, total fish eDNA concentrations, salinity, tide level, and wave height. A gamma distribution was assumed as the error distribution, the “log” function was used as a link function, and the study site and fish species were used as a random effect (i.e., in R, `gamm(abs(IS) ∼ s(explaining_variable), family = Gamma(link=”log”), random = list(site = ∼1, fish_species = ∼1))`, where `s()` indicates a smoothing term). The biological assumption behind this modeling is that explaining variables, such as water temperature, may linearly or nonlinearly influence fish species interactions. Moreover, the effects of explaining variables may randomly vary depending on the study site and fish species. The effect was considered to be statistically clear at *P* < 0.05.

In the second analysis, we focused on the effects of water temperature on the interaction strengths of each fish. In the analysis, GAMM was used again. A gamma distribution was assumed as the error distribution and the “log” function was used as a link function, and the study site was used as a random effect (i.e., in R, `gamm(abs(IS) ∼ s(explaining_variable), family = Gamma(link=”log”), random = list(site = ∼1))`, where `s()` indicates a smoothing term). GAMM was separately performed for each fish species. We also analyzed the effects of other properties on the interaction strengths of each fish species, which are provided in the Supporting Information. Again, the effect was considered to be statistically clear at *P* < 0.05.

### Code and data availability

Source scripts for the analyses and figure generations are archived in Zenodo (https://doi.org/10.5281/zenodo.7865958) and publicly available at Github (https://github.com/ong8181/eDNA-BosoFish-network). DNA data is deposited DDBJ Sequence Read Archive (DRA submission ID = DRA014111).

## Results and Discussion

### Taxonomic diversity and dynamics of fish eDNA

Our multiple MiSeq runs generated 71,892,685 reads in total for the eDNA samples, of which 98% were assigned to fish species, and detected 1,130 molecular operational taxonomic units (MOTUs). We inspected their family-level phylogenies to increase the accuracy of taxonomic assignments, recognizing 856 MOTUs across 33 orders, 167 families, and 466 genera (Table S1). This taxonomic diversity was similar to that of the local fauna (948 species across 33 orders, 158 families, and 493 genera) compiled from a literature survey and museum collections (Table S2). The negative controls produced negligible reads (177 ± 665 reads [mean ± S.D.]), which accounted for *ca*. 0.1% of the positive sample reads. Moreover, all of the reads were assigned to non-target taxa, such as fish species that had never been observed in the study region and freshwater fish species (possibly contaminated from the lab). Therefore, we conclude that any contaminations in our experiments were negligible, and we discarded the sequence reads from the negative control samples.

Sequence reads in the rarefied ASV table were converted to estimated eDNA concentrations by using the eDNA concentrations of a common fish species detected across most samples (Japanese Black Seabream, *A. schlegelii*) (i.e., an analog of the internal spike-in DNA method; see Methods and Fig. S1). The converted ASV table included 23,863 detections and we selected the top 50 most frequently detected species for our time-series analysis. The detection frequencies of these 50 species ranged between 148 and 532 with a mean of 296, and their total detection frequencies reached 14,793 (62.0% of the total detection frequency).

Fish eDNA concentrations and fish species richness showed a clear intra-annual pattern (i.e., seasonality; Fig. 1b, c), which were higher in warmer months (e.g., between July and October) than in colder months (e.g., between January and March) at all sites. Total eDNA concentrations and species richness positively correlated with water temperature (Fig. S2). Seasonal changes in eDNA concentrations were consistent with the patterns of seasonal occurrences in tropical and subtropical fish species in the Boso Peninsula, to which they are transported by the warm Kuroshio Current, settling on the coastal waters during the warmer months and subsequently disappearing during the colder months (Saito, 2019; Senou et al., 2006).

### Reconstruction of the fish species interaction network

We reconstructed fish species interaction networks based on the quantitative, multispecies fish eDNA time series. Regarding the 50 frequently detected fish species, we quantified pairwise information flow between fish species. Figure 2 shows reconstructed fish interaction networks for the 50 frequently detected fish species in the Boso Peninsula. At the regional scale, the linear correlations among water temperature, total eDNA concentrations, interaction strengths, the number of interactions, and species richness was all statistically clear (*P* < 0.05; Fig. S3) except for the linear correlation between water temperature and mean interaction strengths (*P* > 0.05). The interaction strengths became weak as species richness increased, which is consistent with a previous study (Ratzke et al., 2020; Ushio, 2022a), suggesting that understanding the causes and effects of weak interactions is key to understanding the maintenance of species-rich communities.

**Figure 2.**
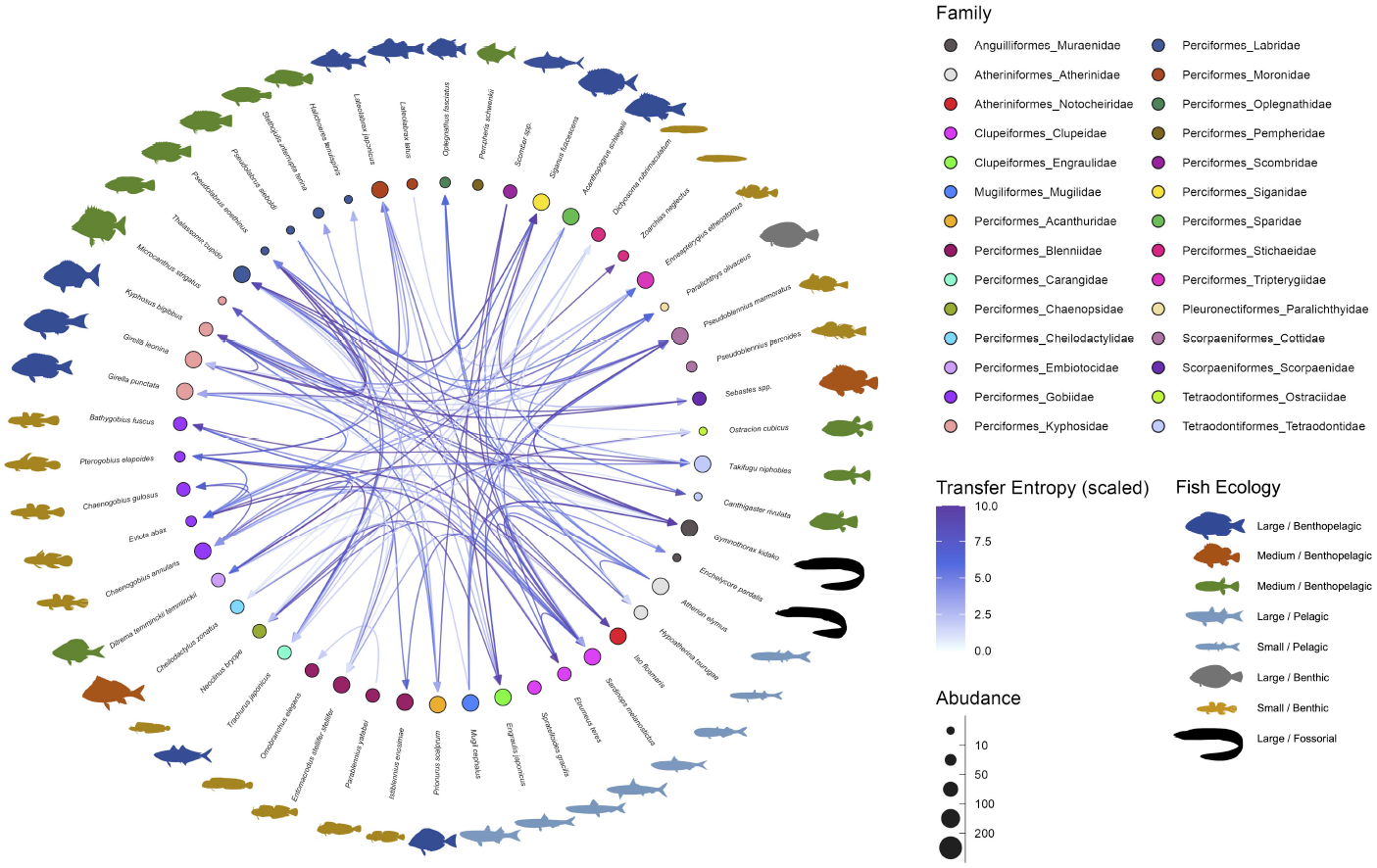
Interaction networks of the fish community in the Boso Peninsula coastal region. The “average” interaction network reconstructed by quantifying information transfer between eDNA time series. Transfer entropy (TE) was quantified by leveraging all eDNA time series from multiple study sites to draw this network. Only information flow larger than 80% quantiles (i.e., strong interaction) was shown as interspecific interactions for visualization. The edge color indicates scaled transfer entropy values, and fish illustration colors represent their ecology (e.g., habitat and feeding behavior). Node colors and node sizes indicate the fish family and fish abundance (total eDNA copy numbers of the fish species), respectively.

Most of the statistically clear information flow may be interpreted from the viewpoint of fish-fish behavior-level interspecific interactions (Table S3 and Supporting Discussion), which is convincing considering the time resolution of our eDNA time series (but see the section “*Potential limitations of the present study*” below). The largest information flow was detected from *Pseudoblennius marmoratus* to *Pseudolabrus eoethinus* (Table S3). These fish species have overlapping habitats, and thus, they may interact. Bidirectional information flow was detected between *P. eoethinus* and *Gymnothorax kidako*. They are both carnivorous fish species and may conduct joint hunting. *Chaenogobius annularis* and *Takifugu niphobles* are often found together in the sand between reefs. *T. niphobles* frequently dive in the sand, and benthos and mysids that are dug up during the dive may be prey for *C. annularis*. Further discussions about the detected information flow are described in Supporting Information (Table S3 and Supplementary Discussion).

### Interaction strengths and environmental variables

We investigated how interaction strengths (i.e., regression coefficients estimated by the MDR S-map) changed with environmental variables (e.g., water temperature) and ecological properties (e.g., species richness and total eDNA concentration) at the community level (Figs. 3 and S4). The in-strengths and out-strengths of fish species interactions were statistically clearly associated with water temperature, species richness, and total eDNA concentration (GAMM, *P* < 0.05; Fig. 3 and Table S4) except for the effects on species richness on the in-strengths (Fig. 3b). In-strengths of the fish-fish interactions increased with water temperature (Fig. 3a), supporting our first hypothesis while out-strengths of the interactions showed an opposite pattern (Fig. 3d), which might suggest there is a difference in the temperature dependence between the in-strengths and out-strengths of the interactions. Indeed, water temperature may influence fish physiological activity (Claireaux et al., 2006; Kishi et al., 2005; Oyugi et al., 2012) often in a complex way, and thus, the community-level influence of water temperature may also be complex as they should arise from the individual-level influence of water temperature on fish. Interaction strengths were also statistically clearly influenced by species richness and total eDNA concentrations (except for Fig. 3b); the interaction strengths decreased with increasing species richness and total eDNA concentration. The effects of salinity, tide and wave were less clear, although the effects were statistically clear except for the effects of tide level on the out-strength (Figs. S4 and S6). Overall, environmental and ecological variables influenced the interaction strengths statistically clearly, but large variations remained unexplained (Figs. S5–S6), suggesting that other factors (e.g., fish ecology, nutrient, and physiological status) may influence the interaction strengths.

**Figure 3.**
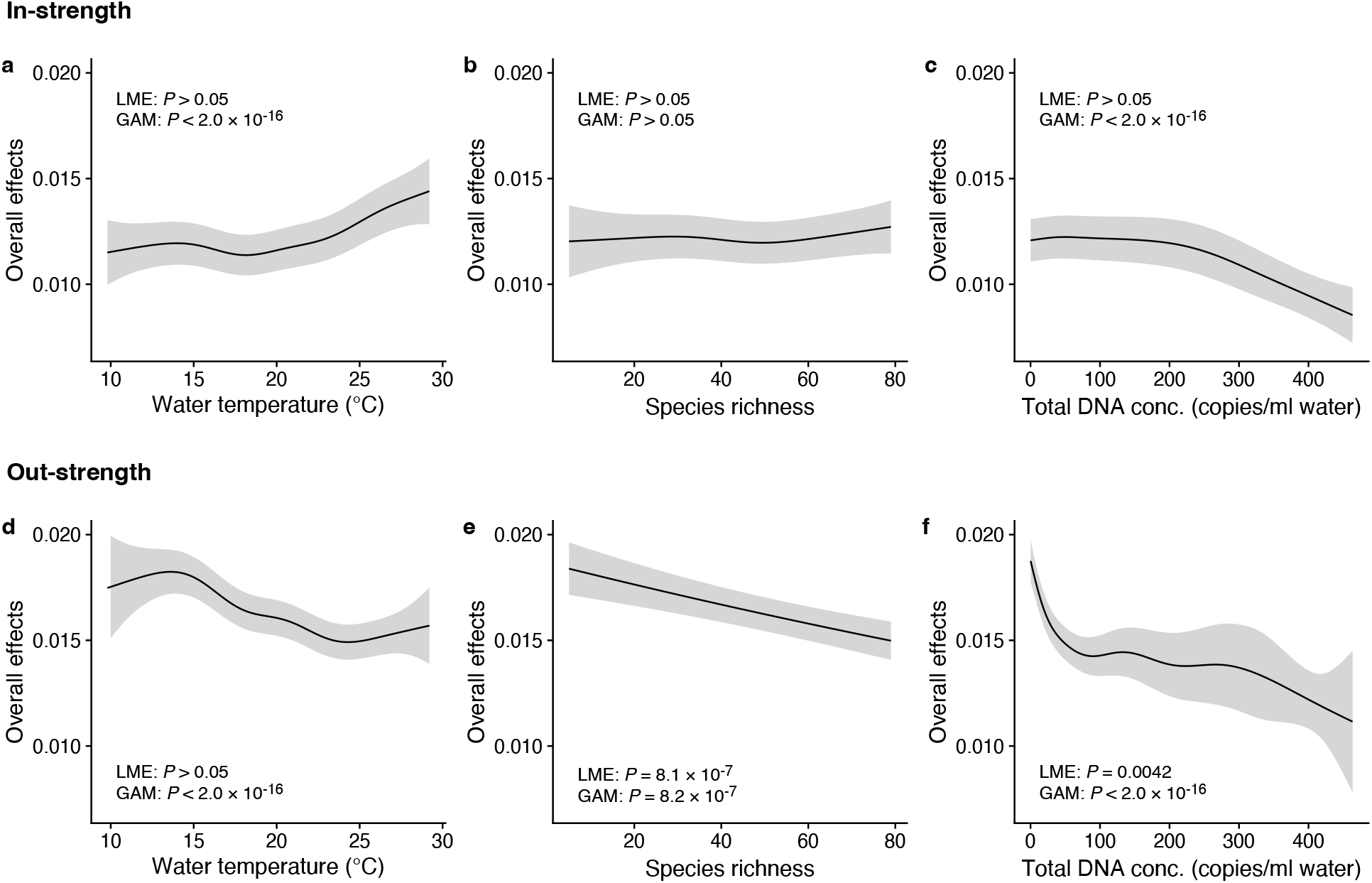
Dependence of interaction strengths on biotic and abiotic variables (50 dominant fish species and 11 study sites were leveraged). The panels show the overall effects of biotic and abiotic variables on interaction strengths of the 50 dominant fish species: Effects of (**a**, **d**) water temperature, (**b**, **e**) species richness, and (**c**, **f**) total eDNA copy numbers. The Y-axis indicates the effects of the variables on fish-fish interaction strengths quantified by the MDR S-map method. **a**–**c** show the effects on the species interactions that a focal species receives (i.e., In-strength), and **d**–**f** show the effects on the species interactions that a focal species gives (i.e., Out-strength). The line indicates the average effects estimated by the general additive model (GAM), and the grey region indicates 95% confidential intervals. LME and GAM indicate the statistical clarity of the linear mixed model portion and GAM portion, respectively. Detailed statistical results and raw data are shown in Table S4 and Fig. S5, respectively.

We also investigated how temperature influenced the interaction strength of individual fish species. Figure 4 shows how interaction strengths among fish species changed with water temperature. For visualization purpose, we show fish species of which interaction strengths changed with water temperature highly statistically clearly (*P* < 0.0001). The in-strengths of several fish species clearly increased at higher water temperatures: for example, *Halichoeres tenuispinis*, *Macrocanthus strigatus*, *Pempheris schwenkii*, *Stethojulis interrupta terina*, and *Thalassoma cupido* (Fig. 4a). On the other hand, in-strengths of some fish species and out-strengths decreased at higher water temperatures: for example, in-strengths of *Ditrema temminckii temminckii*, *Engraulis japonicus*, and *Girella punctata*, and out-strengths of five fish species (Fig. 4a,b). These results support our second hypothesis that the relationship between the patterns of temperature and interaction strengths varies among fish species. These results suggest that, although water temperature may have strong influences on fish-fish interaction strengths in general, the sign of temperature effect (i.e., positive or negative) can vary depending on fish species and environmental conditions. Previous studies showed that fish physiological activities, such as feeding rates, growth rates, and swimming speed, were influenced by water temperature (Claireaux et al., 2006; Kishi et al., 2005; Oyugi et al., 2012). In addition, the direction (i.e., positive or negative) of the temperature effects on fish metabolic activities may be species-specific (Oyugi et al., 2012), and this species specificity may underlie the species-specific effects of temperature on the interaction strength.

**Figure 4.**
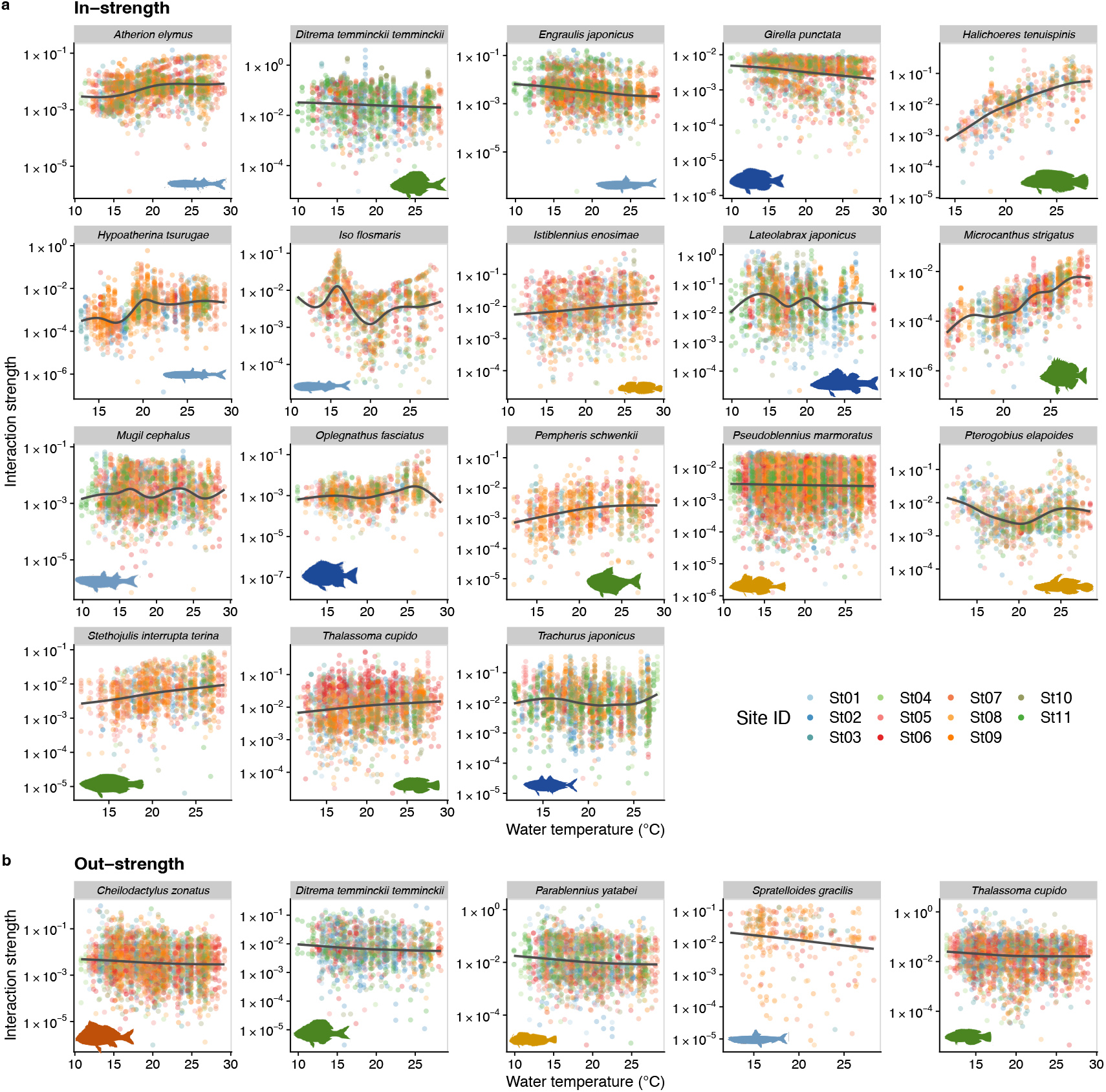
Temperature dependence of fish species interactions at the species level. **a** and **b** show temperature effects on fish species interactions quantified by the MDR S-map method. Note that the MDR S-map enables quantifications of interaction strengths at each time point, and thus the number of data points is large. (**a**) Points indicate the species interactions that a focal species (indicated by the strip label and fish image) receives (i.e., In-strength). (**b**) Points indicate the species interactions that a focal species (indicated by the strip label and fish image) gives (i.e., Out-strength). For **a** and **b**, only fish species of which interactions are statistically clearly affected by water temperature are shown (to exclude fish species with relatively weak temperature effects, *P* < 0.0001 was used as a criterion here). Point color indicates the study site. Gray line is drawn by GAM (the study sites were averaged for visualization purpose).

### Potential limitations of the present study and future perspectives

The present results showing that the strengths of fish-fish interactions depend on water temperature rely on several assumptions that were not fully investigated, and thus, careful interpretations and discussions are required. First, the extent to which fish eDNA concentrations accurately represent fish abundance is an open question. Previous studies demonstrated that the quantity of eDNA may be a proxy of fish abundance and/or biomass (i.e., a positive correlation between the quantity of eDNA and fish abundance/biomass) (Takahara et al., 2012; Thomsen et al., 2012; Yamamoto et al., 2016). Our nonlinear time series analysis used the information embedded in fluctuations in time series, and time series are always standardized before the analysis. Therefore, species-specific differences in the copy numbers of the genetic marker and eDNA release rates, which may bias the estimation of absolute abundance, do not cause serious biases in the reconstruction of the networks and estimations of interaction strengths. However, accurate estimations of the absolute abundance of fish will improve the accuracy of analyses, and the integration of eDNA concentration data, knowledge on eDNA dynamics (i.e., release and degradation rates), and the hydrodynamic modeling of ocean water flow will be a promising approach (e.g., as in Fukaya et al., 2021).

Another limitation is that the reconstructed interaction network was not fully validated in the present study. Although previous studies demonstrated that network reconstructions based on a time series analysis work reasonably well and show meaningful patterns (Runge et al., 2019; Sugihara et al., 2012; Ushio, 2022a; Ushio et al., 2018a), experiment/observation-based validations of the reconstructed network have not yet been performed. Potential causal interactions between fish species may be reasonably interpreted (Supplementary Discussion); however, developing a framework to efficiently validate “previously unknown” interactions detected by eDNA data and a nonlinear time series analysis is an important future direction.

Third, although we showed that temperature effects on interaction strength are statistically clear and common in the Boso peninsula coastal region, the range of water temperature in the present analysis was still not broad and there is currently no evidence that we can generalize these results to other regions. Nonetheless, the advantages of the eDNA method are the low cost and minimal time and labor needed for field sampling as well as the potential scalability of the library preparation process (Ushio et al., 2022). Therefore, frequent (Ushio, 2022a) and large spatial-scale ecological monitoring (e.g., ANEMONE DB; https://db.anemone.bio/) is possible, which will provide a more detailed understanding of the temperature-interaction strength relationships of fish on a larger spatial scale.

Lastly, other environmental conditions may covary with temperature and influence fish-fish interactions in a species-specific way (Figs. S7 and S8). Although the effect of temperature on the interaction strengths was the strongest at the community level among the environmental variables (Table S4), there is a possibility that the temperature effect we observed might not be direct. For example, water temperature and oxygen concentration covary and both directly/indirectly affect fish physiology (Salvatteci et al., 2022). Because direct field manipulations to validate our results are challenging, robust conclusions about the temperature-sensitivity of fish interactions may only be made by integrating results from different approaches (e.g., small-scale lab experiment- and time series-based causal analysis). In nature, multiple environmental variables are continuously changing, and the interaction strengths may fluctuate through time and space being affected by the environmental variables, as shown in previous studies (Ushio, 2022a; Ushio et al., 2018a). Conversely, interactions detected and quantified under a controlled environment might not necessarily be observed under field conditions. Our research framework that enables the detection and quantification of interactions in nature provides a complementary view about fish-fish interactions, which would play a critical role in understanding the effects of temperature on fish-fish interactions.

### Implications for fish community assembly and the effect of global climate change

Water temperature has significant influences on marine community composition and diversity at the global spatial scale and historical time scale (Tittensor et al., 2010; Yasuhara and Deutsch, 2023), but the mechanism of temperature effects on fish community assembly is not fully understood. Recent studies have shown that temperature (and oxygen) plays an important role in determining fish body size at the historical time scale (Salvatteci et al., 2022; Yasuhara and Deutsch, 2022). As fish body size plays a critical role in interspecific interactions such as predator-prey interactions (e.g., predator-prey mass ratio; Nakazawa et al., 2011), temperature-induced body size changes may influence interspecific interactions at a longer time scale. On the other hand, our study showed that temperature may induce changes in fish-fish interactions at a relatively short time scale (weeks to months), perhaps via temperature effects on the physiological activity of fish individuals. These suggest that temperature effects on fish community assembly involve effects at different time scales, and thus, integrating results from different temporal (and spatial) scales are necessary to understand fish community assembly processes in nature.

In addition, our study revealed that temperature effects on fish-fish interactions depend on fish species identity. This suggests that, even in the same habitat, the temperature-sensitivity of fish-fish interactions is variable and fish species-specific, and that consequences of changing temperature in the community assembly process may be complex. For example, increased water temperature strengthens interactions received by *Halichoeres tenuispinis* and *Microcanthus stringatus* (Fig. 4), which may destabilize the population dynamics of these species. In contract, increased water temperature may exert the opposite effects on *Engraulis japonicus* and *Girella punctata* (Fig. 4), that is, weakened interactions and stabilized population dynamics. How these varying responses to temperature change, or “response diversity” (Ross et al., 2023), influences overall community dynamics remains unclear. Our study provides a practical framework to quantify response diversity using time series (the S-map method calculates the first derivative as an interaction strength, and it is an analog of the additive model-based method proposed by Ross et al. 2023), and quantifying species-specific responses to environmental changes and their diversity would be key to predicting the community-level responses under climate change.

### Conclusions

The present study demonstrated that the strengths of fish species interactions changed with water temperature under field conditions. For several fish species, species interactions were intensified in warmer water and for some other fish species, species interactions were weakened in warmer water. This may change the correlation and distribution of interaction strengths in a community, which may consequently influence community dynamics and stability in a complex way (Allesina et al., 2015; Tang et al., 2014; Ushio et al., 2018a). Therefore, a more detailed understanding of the effects of environmental conditions on species interactions under field conditions will be required to improve our capability to understand, forecast, and even manage natural ecological communities and dynamics, which is particularly important under ongoing global climate change. Developments and improvements in eDNA techniques and statistical analyses are still required; however, because eDNA analysis is potentially applicable to any type of organisms even if they are difficult to be detected using traditional methods, our framework integrating an eDNA analysis and advanced statistical analyses pave a way to understand and forecast dynamics of various ecological communities under field conditions.

## Acknowledgments

We thank T. Komai, R.O. Gotoh, T. Sunobe, and K. Takiguchi for assisting with the bimonthly collection of eDNA samples. This research was supported by JSPS KAKENHI (B) Grant Number 20H03323, the Hakubi Project in Kyoto University, and The Hong Kong University of Science and Technology Startup Funding to MU, and JSPS KAKENHI (B) Grant Numbers JP19H03291, JP22H02691, and MEXT OGAP Project Grant Number JPMXD0618068274 to MM.

## Author contributions

All authors agreed to the submission. MU and MM conceived the idea and designed the research; MM, TS, and TF collected the samples, performed eDNA metabarcoding and analyzed sequence data; SS and RM performed quantitative PCR analysis; MU compiled data; MU and YO analyzed data; MU and MM wrote the first draft and completed the final manuscript with contributions from all coauthors.

