## Supplementary Figures S1-S8 for "Temperature sensitivity of the interspecific interaction strength of coastal marine fish communities"

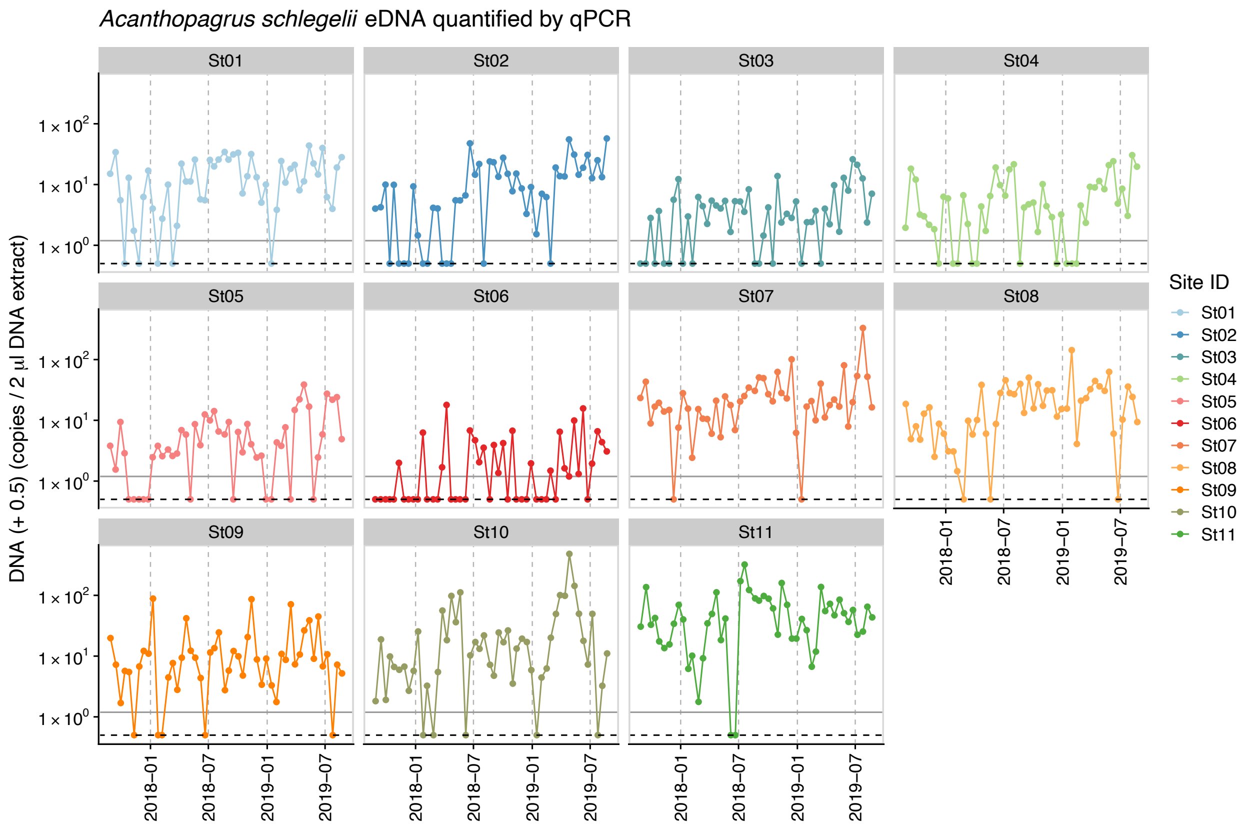


**Figure S1 | Dynamics of environmental DNA (eDNA) copy numbers of Japanese black seabream (*Acanthopagrus schlegelii*) that was used as an internal standard.** Gray horizontal line indicates the minimum eDNA copy number of Japanese black seabream detected. Approximately 83.3% of samples contain detectable concentrations of the standard DNA, which was used to convert sequence reads to eDNA copy numbers. For samples which do not contain the standard DNA, we assume that it contains the minimum amount of eDNA copy numbers (i.e., 0.69 copies / 2 $\mu$l extracted DNA; gray horizontal line).

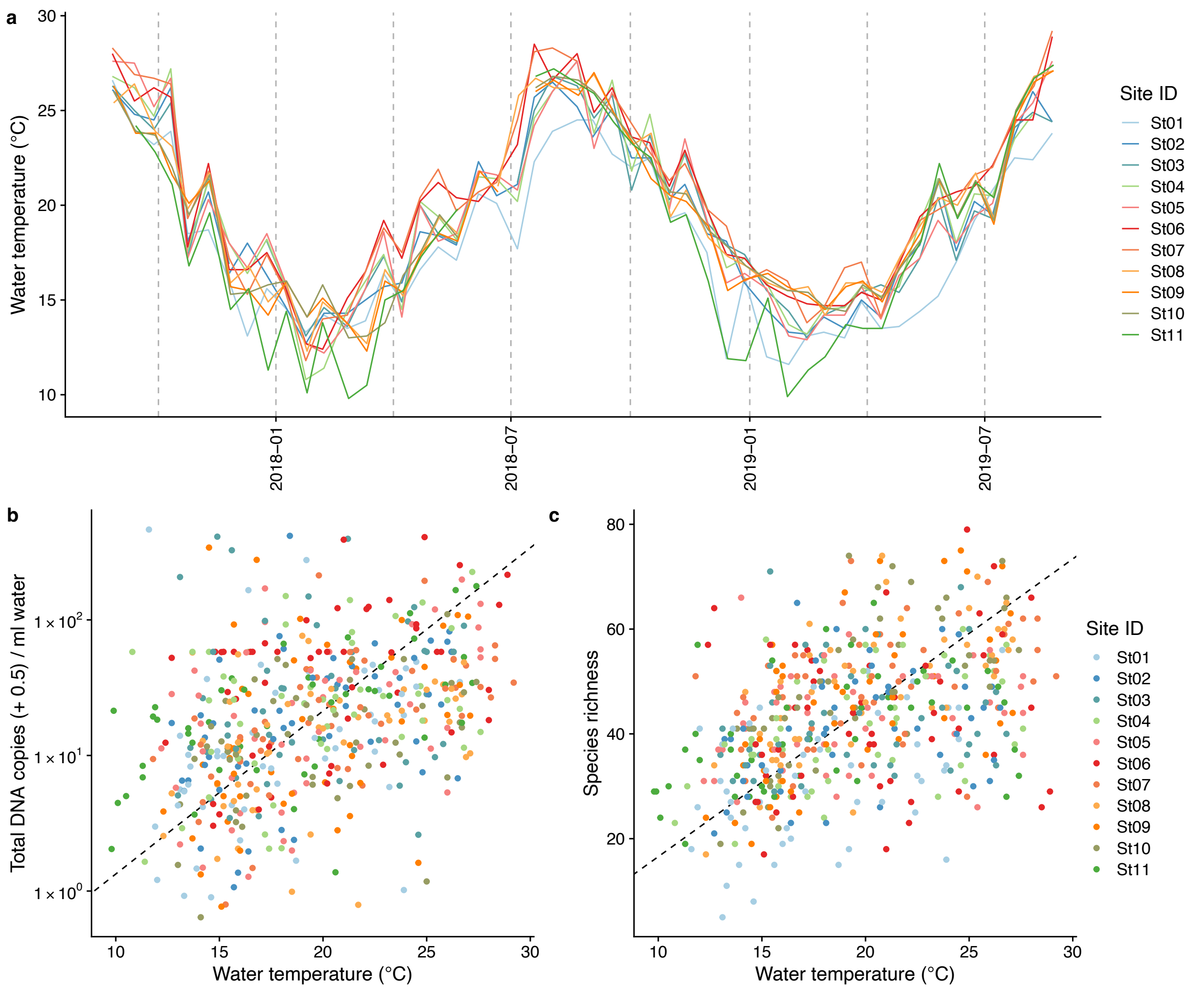


**Figure S2 | Dynamics of water temperature and the relationships between water temperature and total eDNA concentration and fish species richness.** (**a**) Dynamics of sea surface water temperature at the sampling sites. (**b**) The relationship between water temperature and total eDNA copy numbers, and (**c**) the relationship between water temperature and fish species richness. Colors indicate sampling sites. For **b** and **c**, dashed lines indicate standardized major axis regression.

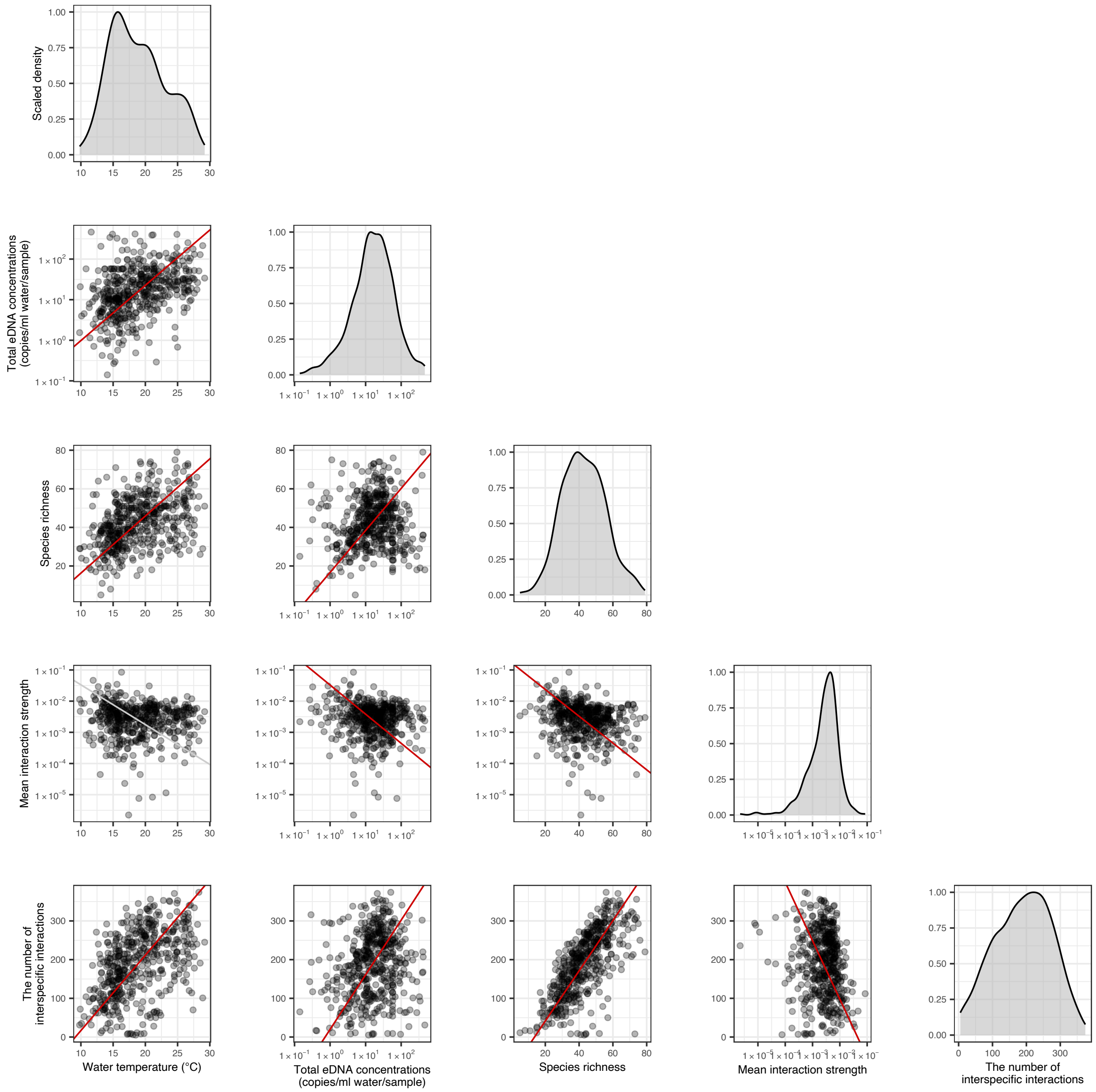


**Figure S3 | The relationships between network properties and environmental variables**. Diagonal panels show density distributions of the data, and lower triangle area shows scattered plots. Each point represent one water sample at each study site, meaning that "Mean interaction strength" is a mean value at the community level. Solid red and gray lines indicate statistically clear ($P < 0.05$) and unclear ($P > 0.05$) standardized major axis regression, respectively.

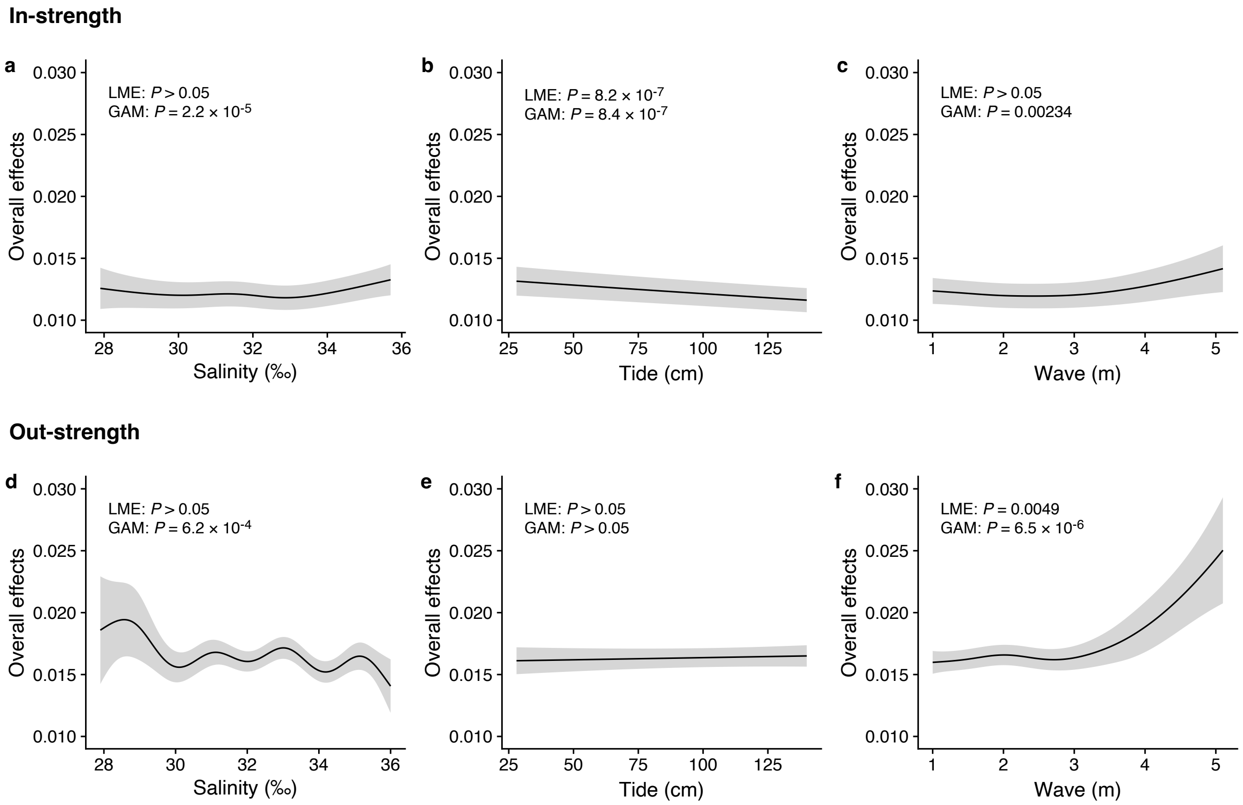


**Figure S4 | Dependence of interaction strengths on additional abiotic variables**. The panels show the overall effects of additional abiotic variables on interaction strengths of the 50 dominant fish species: Effects of (**a**, **d**) salinity, (**b**, **e**) tide level (cm), and (**d**, **f**) wave (m). The $y$-axis indicates the effects of environmental variables on fish-fish interaction strengths quantified by the MDR S-map method. **a**-**c** show the effects on the species interactions that a focal species receives (i.e., In-strength), and **d**-**f** show the effects on the species interactions that a focal species gives (i.e., Out-strength). The line indicates the average effects estimated by the general additive model (GAM), and the grey region indicates 95% confidential intervals. LME and GAM indicate the statistical clarity of the linear mixed model portion and GAM portion, respectively. Detailed statistical results are shown in Table S4.

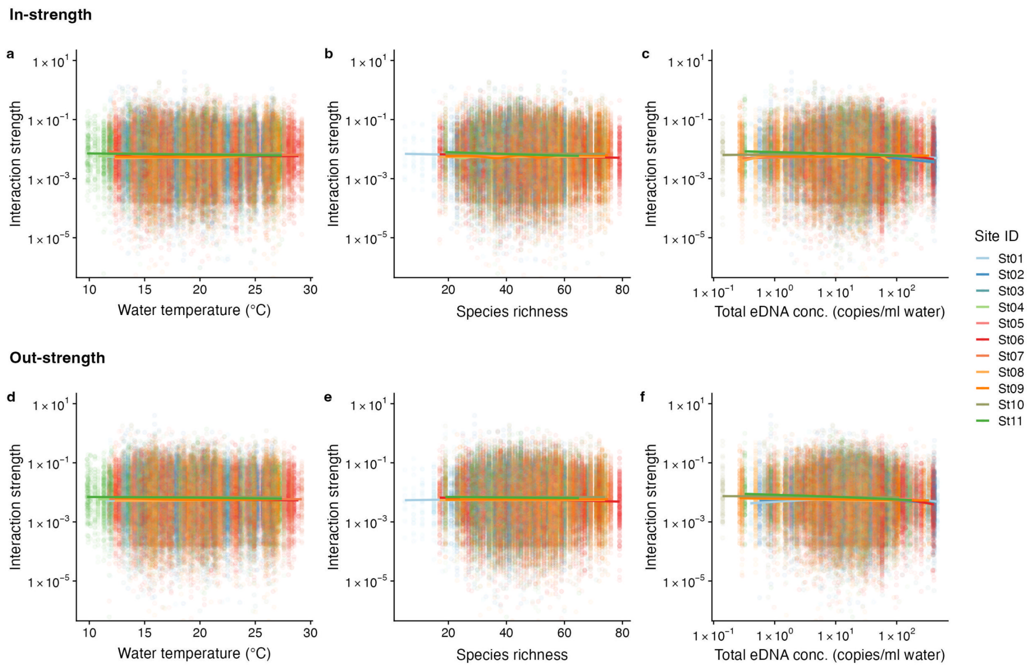


**Figure S5 | The relationship between interaction strengths and water temperature, species richness, and total DNA concentrations**. The panel shows the overall relationship between interaction strengths of the 50 dominant fish species and (**a**, **d**) water temperature, (**b**, **e**) species richness, and (**d**, **f**) total DNA concentration. The $y$-axis indicates the interaction strength between fish species quantified by the MDR S-map method. Note that the MDR S-map enables quantification of interaction strengths at each time point, and thus the number of data points is large (also true in Figs. S6-S8). (**a**-**c**) Points indicate the species interactions that a focal species receives (i.e., In-strength), and (**d**-**f**) points indicate the species interactions that a focal species gives (i.e., Out-strength). The line indicates the nonlinear regression line estimated by the general additive model. Colors of points and lines indicate the study site.

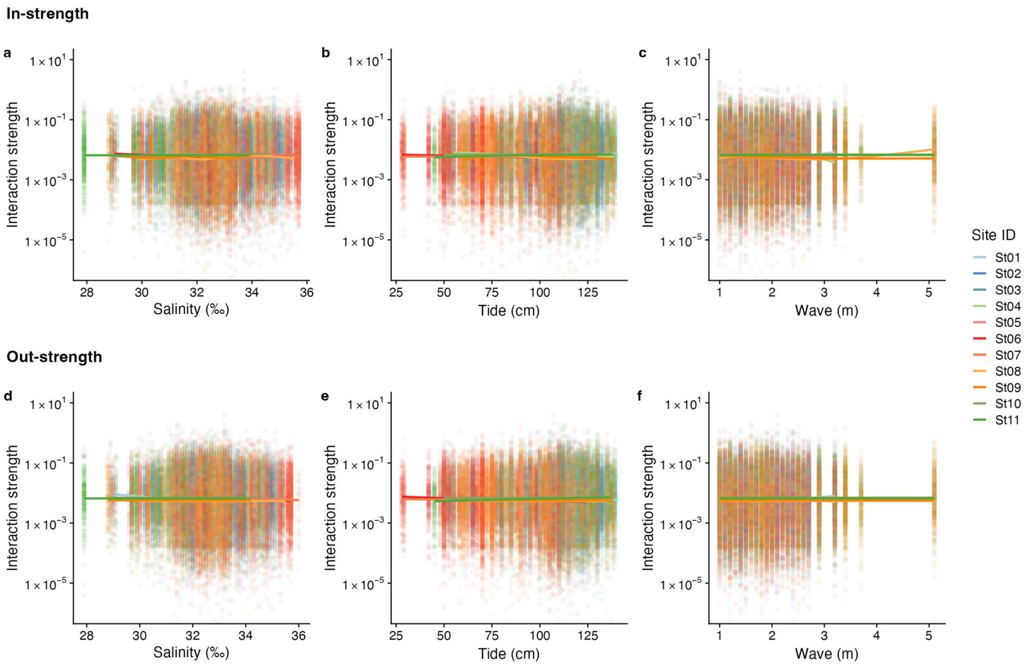


**Figure S6 | The relationship between interaction strengths and salinity, tide level, and wave**. The panel shows the overall relationship between interaction strengths of the 50 dominant fish species and (**a**, **d**) salinity, (**b**, **e**) tide level (cm), and (**c**, **f**) wave (m). The $y$-axis indicates the interaction strength between fish species quantified by the MDR S-map method. Note that the MDR S-map enables quantifications of interaction strengths at each time point, and thus the number of data points is large. (**a**-**c**) Points indicate the species interactions that a focal species receives (i.e., In-strength), and (**d**-**f**) points indicate the species interactions that a focal species gives (i.e., Out-strength). The line indicates the nonlinear regression line estimated by the general additive model. Colors of points and lines indicate the study site.

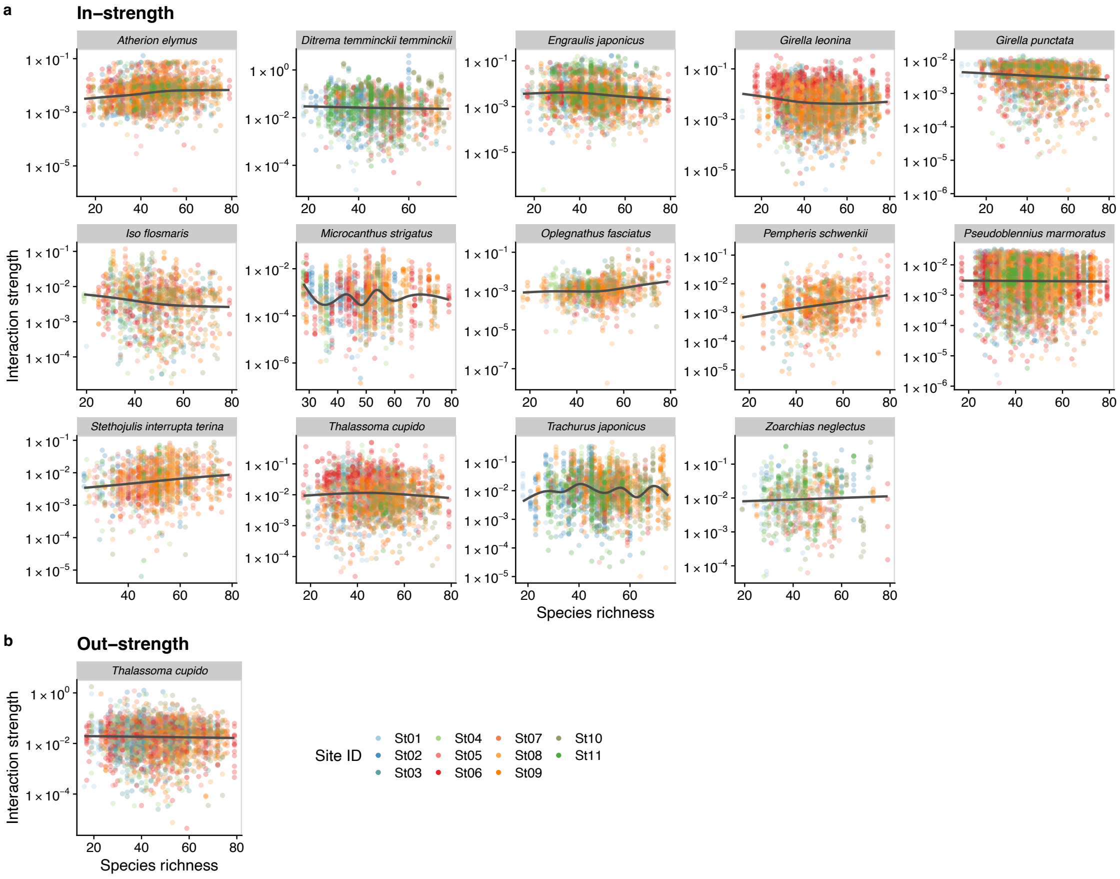


**Figure S7 | Dependence of fish species interactions on species richness at the fish species level**. **a** and **b** show effects of species richness on fish species interactions quantified by the MDR S-map method. (**a**) Points indicate the species interactions that a focal species (indicated by the strip label and fish image) receives (i.e., In-strength). (**b**) Points indicate the species interactions that a focal species (indicated by the strip label and fish image) gives (i.e., Out-strength). For **a** and **b**, only fish species of which interactions are statistically clearly affected by water temperature are shown (to exclude fish species with relatively weak temperature effects, $P < 0.0001$ was used as a criteria here). Point color indicates the study site. Gray line is drawn by GAM (the study sites were averaged for visualization purpose).

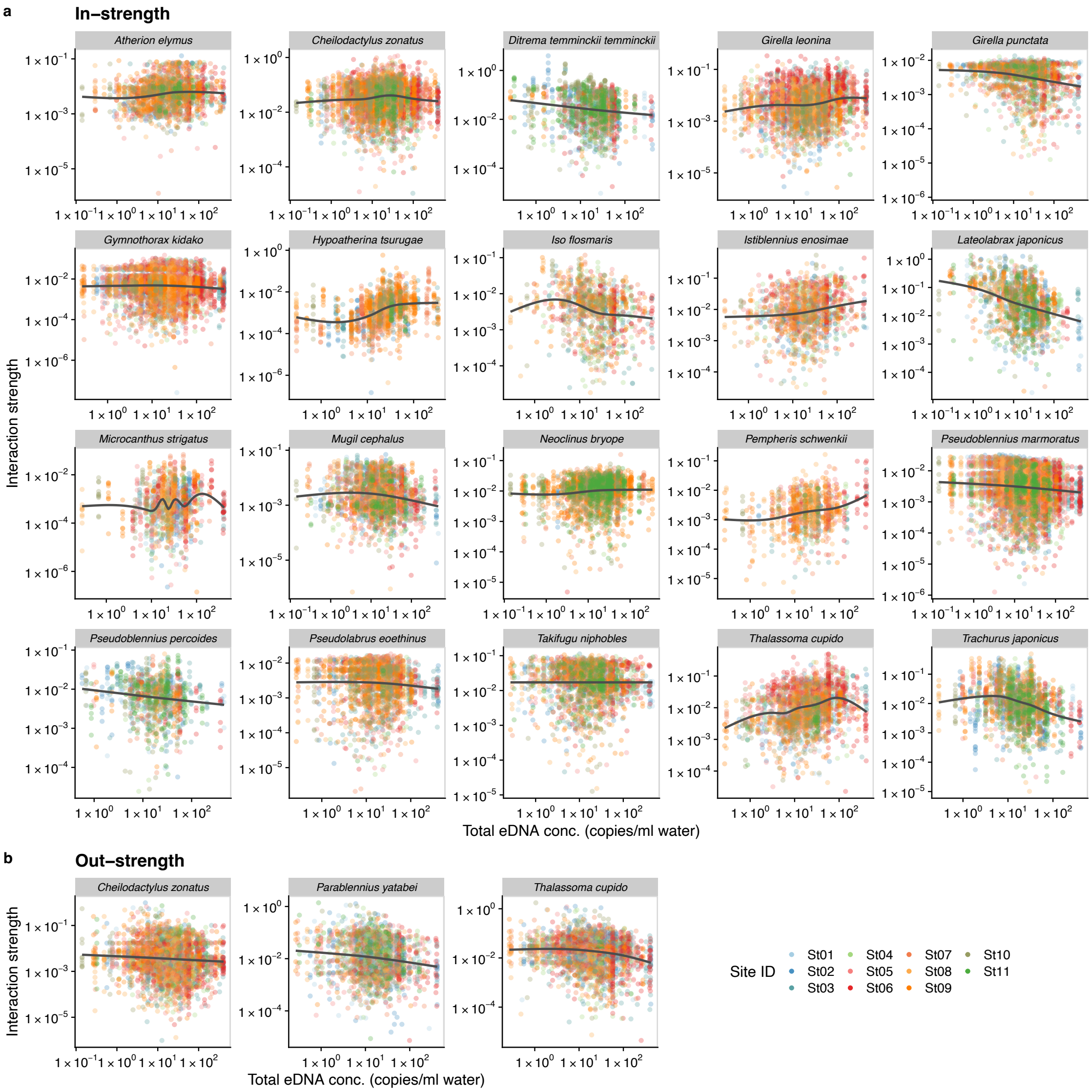


**Figure S8 | Dependence of fish species interactions on the total DNA concentration (an index of total fish abundance) at the species level**. **a** and **b** show effects of the total DNA concentrations on fish species interactions quantified by the MDR S-map method. (**a**) Points indicate the species interactions that a focal species (indicated by the strip label and fish image) receives (i.e., In-strength). (**b**) Points indicate the species interactions that a focal species (indicated by the strip label and fish image) gives (i.e., Out-strength). For **a** and **b**, only fish species of which interactions are statistically clearly affected by water temperature are shown (to exclude fish species with relatively weak temperature effects, $P < 0.0001$ was used as a criteria here). Point color indicates the study site. Gray line is drawn by GAM (the study sites were averaged for visualization purpose).
