## Supplementary Text for "Temperature sensitivity of the interspecific interaction strength of coastal marine fish communities"

**Contents:**

**Supplementary Materials and Methods** | Detailed descriptions of materials and methods.
**Supplementary Discussion** | Interpretations of and discussions on detected interactions
**Figure S1** | Dynamics of environmental DNA (eDNA) copy numbers of Japanese black seabream (*Acanthopagrus schlegelii*) that was used as an internal standard.
**Figure S2** | Dynamics of water temperature and relationships between water temperature, total eDNA concentration, and fish species richness.
**Figure S3** | The relationships between network properties and environmental variables.
**Figure S4** | Dependence of interaction strengths on additional abiotic variables.
**Figure S5** | The relationship between interaction strengths and water temperature, species richness, and total DNA concentration.
**Figure S6** | The relationship between interaction strengths and salinity, tide level, and wave height.
**Figure S7** | Dependence of fish species interactions on species richness at the species level.
**Figure S8** | Dependence of fish species interactions on the total DNA concentration (an index of total fish abundance) at the species level.
**Table S1**| List of species detected from the eDNA metabarcoding with raw numbers of reads.
**Table S2**| A faunal inventory of coastal marine fishes of Chiba prefecture compiled from museum collections and literature surveys.
**Table S3** | Top 20 fish-fish interactions based on UIC.
**Table S4** | Results of GAMM between interaction strengths and environmental and ecological properties.

#### Supplementary Materials and Methods

##### *Seawater sampling and on-site filtration*

We employed low-tech bucket sampling to collect seawater using a folding 7.8-L polypropylene bucket (Soft Bucket 8, ISETO, Osaka, Japan) fastened to a 15-m rope (vinylon rope, φ6 mm). Prior to seawater sampling, we wore disposable gloves on both hands and assembled a set of on-site filtration kits consisting of a Sterivex filter cartridge (pore size of 0.45 µm; Merck Millipore, MA, USA) and a 50-ml disposable syringe with a Luer lock connector (TERUMO, Tokyo, Japan). We then thoroughly decontaminated the bucket with a foam-style 10% bleach solution and brought the equipment to the sampling site. We fixed the end of the 15-m rope fastened to the bucket and collected surface seawater by casting and retrieving the bucket full of seawater. We repeated this collection of seawater ten times to minimize sampling biases at each station.

Two researchers performed on-site filtration using two pairs of the above kit (filter cartridge + syringe) to obtain duplicate samples. With each collection of seawater, we removed the filter cartridge from the syringe, drew approximately 50-mL seawater into the syringe by pulling the plunger, reattached the filter cartridge to the syringe, and pushed the plunger for the filtration of seawater. We repeated this step twice in a single cast of the bucket, and the final filtration volume reached 1,000 mL × 2 with ten casts of the bucket. When the filter was clogged before reaching 1,000-mL filtration, we recorded the total volume of water filtered. After on-site filtration, we added 1.6 mL of RNAlater (Thermo Fisher Scientific, DE, USA) to the cartridge to prevent eDNA degradation. We made a filtration blank (FB) by filtering 500 mL of purified water in the same manner at the end of each day of water sampling. We transported the filtered cartridges to the laboratory in a portable cooler with ice packs and kept these cartridges at –20˚C in the freezer until eDNA extraction.

##### *eDNA extraction*

We thoroughly sterilized the workspace and equipment before DNA extraction. We used filtered pipette tips and conducted all eDNA extraction manipulations in a dedicated room that was separate from the pre- and post-PCR rooms to safeguard against cross-contamination from PCR products.

We extracted eDNA from the filter cartridges using the DNeasy Blood & Tissue kit (Qiagen, Hilden, Germany) following the methods developed and visualized by Miya et al. (2016) with slight modifications (Minamoto et al., 2021; Miya and Sado, 2019a). Briefly, we connected the inlet port of each filter cartridge to a 2.0-mL collection tube and tightly sealed the connection between the cartridge and collection tube with Parafilm. We inserted the combined unit into a 15-mL conical tube and centrifuged the capped conical tube at 6,000 × *g* for 1 min to remove redundant seawater and RNAlater. After centrifugation, we discarded the collection tube and used an aspirator (QIAvac 24 Plus, Qiagen, Hilden, Germany) to completely remove any liquid remaining in the cartridge.

We subjected the filter cartridge to lysis using proteinase K. Before lysis, we mixed PBS (220 µL), proteinase K (20 µL), and buffer AL (200 µL), and gently pipetted the mixed solution into the cartridge and incubated the cartridge at 56˚C for 20 min while stirring the cartridge using a rotator (10 rpm; Mini Rotator ACR-100, AS ONE, Tokyo, Japan). After the incubation, we collected the lysate and purified the DNA extract (*ca*. 900 µL) using the DNeasy Blood and Tissue kit following the manufacturer’s protocol and set the final elution volume at 200 μL. We also made an extraction blank (EB) during this process in addition to FB.

##### *Paired-end library preparation and MiSeq sequencing*

We thoroughly sterilized the workspace and equipment in the pre-PCR area before library preparation. We used filtered pipette tips while performing pre- and post-PCR manipulations in two different dedicated rooms to safeguard against cross-contamination.

We employed two-step PCR for paired-end library preparation using the MiSeq platform (Illumina, CA, USA). We generally followed the methods developed by Miya et al. (2015) and subsequently modified by Miya & Sado (2019b). In the first-round PCR (1st PCR), we used a mixture of the following six primers: MiFish-U-forward, MiFish-U-reverse, MiFish-E-forward-v2, MiFish-E-reverse-v2, MiFish-U2-forward, and MiFish-U2-reverse (Table 1). These primer pairs amplified a hypervariable region of the mitochondrial 12S rRNA gene (*ca*. 172 bp; hereafter called the "MiFish sequence") and appended primer-binding sites (5′ ends of the sequences before six random bases [Ns]) for sequencing at both ends of the amplicon. Six Ns in the middle of these primers enhanced cluster separation on the flow cells during initial base-call calibrations on the MiSeq platform.

The 1st PCR consisted of 35 cycles with a 12-μL reaction volume containing 6.0 μL of 2 × KAPA HiFi HotStart ReadyMix (KAPA Biosystems, MA, USA), 2.8 μl of a mixture of the three MiFish primer sets at a volume ratio of 2:1:1 (U:E:U2 forward and reverse primers; 5 μM), 1.2 μL of sterile distilled $H_2O$, and 2.0 μL of the eDNA template. To minimize PCR dropouts during the 1st PCR (Doi et al., 2019; Miya et al., 2020), we performed eight technical replicates for the same eDNA template using a strip of eight tubes (200 μL). The thermal cycle profile after an initial 3-min denaturation at 95˚C was as follows: denaturation at 98˚C for 20 s, annealing at 65˚C for 15 s, and extension at 72˚C for 15 s, with a final extension at the same temperature for 5 min. We also prepared a 1st PCR blank (1B) during this process, in addition to FB and EB. However, we did not perform eight replications and only used a single tube for each of the three blanks (FB, EB, and 1B) to minimize costs.

After completing the 1st PCR, we pooled an equal volume of PCR products from each of the eight replicates in a single 1.5-mL tube. We purified pooled products using a GeneRead Size Selection kit (Qiagen, Hilden, Germany) following the manufacturer's GeneRead DNA Library Prep I Kit protocol. The column purification process was repeated twice. We subsequently quantified the purified target products (*ca*. 300 bp) using TapeStation 2200 (Agilent Technologies, Tokyo, Japan), diluted to 0.1 ng/μL using Milli Q water, and used the diluted products as templates for the second-round PCR (2nd PCR). Regarding the three blanks (FB, EB, and 1B), we purified the 1st PCR products in the same manner, but did not quantify the purified PCR products. We instead diluted them according to an average dilution ratio for positive samples, following which we used the diluted products as templates for the 2nd PCR.

In the 2nd PCR, we used the two primers to append dual-indexed sequences (8 nucleotides indicated by **X**) and flow cell-binding sites for the MiSeq platform: 2nd PCR-forward and 2nd PCR-reverse (Table 1). We performed the 2nd PCR with 10 cycles of a 15-μL reaction volume containing 7.5 μL of 2 × KAPA HiFi HotStart ReadyMix, 0.9 μL of each primer (5 μM), 3.9 μL of distilled $H_2O$, and 1.9 μL of the template (0.1 ng/μL with the exceptions of the three blanks). The thermal cycle profile after an initial 3-min denaturation at 95˚C was as follows: denaturation at 98˚C for 20 sec, annealing and extension combined at 72˚C (shuttle PCR) for 15 sec with the final extension at the same temperature for 5 min. We also made a 2nd PCR blank (2B) during this process in addition to FB, EB, and 1B. In total, we made 195 blanks (FB = 60, EB = 50, 1B = 50, 2B = 35) and subjected them to the above library preparation procedure to monitor contamination during the on-site filtration, subsequent DNA extraction, and 1st and 2nd PCR of the 550 samples.

We adjusted the number of samples per MiSeq sequencing to obtain approximately 100,000 reads per sample. We pooled each of the individual paired-end libraries in an equal volume into a 1.5-mL tube. We then electrophoresed the pooled libraries using a 2% E-Gel Size Select agarose gel (Invitrogen, CA, USA) and excised the target amplicons (*ca*. 370 bp). The concentrations of the size-selected libraries were measured using a Qubit dsDNA HS assay kit and Qubit fluorometer (Life Technologies, CA, USA), diluted to 10–12.0 pM with HT1 buffer (Illumina, CA, USA), and sequenced on the MiSeq platform using a MiSeq v2 Reagent Kit for 2 × 150 bp PE (Illumina, CA, USA) with a PhiX Control library (v3) spike-in (expected at 5%) following the manufacturer's protocol. All raw DNA sequence data and associated information are deposited in DDBJ/EMBL/GenBank and are available from the accession number DRA014111.

##### *Sequence data preprocessing*

We performed data preprocessing and analyses of raw MiSeq reads from the MiSeq run using PMiFish ver. 2.4 (https://github.com/rogotoh/PMiFish; Miya et al. 2020) according to the following steps:

1) Forward (R1) and reverse (R2) reads were merged by aligning the two reads using the *fastq merge pairs* command. During this process, short reads (<100 bp) after tail trimming and paired reads with too many differences (>5 positions) in the aligned region (*ca*. 65 bp) were discarded; 2) primer sequences were removed from merged reads using the *fastx truncate* command; 3) reads without primer sequences underwent quality filtering using the *fastq filter* command to remove low quality reads with an expected error rate of >1% and short reads of <20 bp; 4) preprocessed reads were dereplicated using the *fastx uniques* command and all singletons, doubletons, and tripletons were removed from subsequent analyses to avoid false positives following the recommendation by the author of the program (Edgar, 2010); 5) dereplicated reads without singletons, doubletons, and tripletons were denoised using the *unoise3* command to generate amplicon sequence variants (ASVs) (Edgar, 2016); 6) ASVs were rarefied to the approximate minimum read number (20,000); and 7) ASVs were subjected to taxon assignments to species names (molecular operational taxonomic units; MOTUs) using the *usearch global* command with a sequence identity of >98.5% with the reference sequences (two nucleotide differences allowed) and a query coverage of ≥90%.

##### *Taxon assignment*

ASVs with sequence identities of 80–98.5% were tentatively assigned "U98.5" labels before the corresponding species names with the highest identities (e.g., U98.5 *Pagrus major*) and were subjected to clustering at the level of 0.985 using the *cluster smallmem* command. An incomplete reference database necessitates this clustering step, which enables the detection of multiple MOTUs for identical species names. Multiple MOTUs were annotated as "gotu1, 2, 3…" and all outputs (MOTUs plus U98.5 MOTUs) were tabulated with read abundance. ASVs with sequence identities of <80% (saved as "no hit") were excluded from the above taxon assignments and downstream analyses because they were all non-fish organisms. MiFish DB ver. 43 was used for taxon assignment, comprising 7,973 species distributed across 464 families and 2675 genera.

To refine the above taxon assignments, family-level phylogenies were reproduced from

MiFish sequences from MOTUs, U98.5 MOTUs, and reference sequences (contained in the MiFish DB ver. 43) belonging to these families. In each family, representative sequences (most abundant reads) from MOTUs and U98.5 MOTUs were assembled, and all reference sequences were added from that family and saved in the FASTA format. Combined FASTA-formatted sequences were subjected to multiple alignments using MAFFT 7 (Katoh and Standley, 2013) with a default set of parameters. A neighbor-joining (NJ) tree was subsequently constructed with the aligned sequences in MEGA X (Stecher et al., 2020) using Kimura two-parameter distances. Distances were calculated using the pairwise deletion of gaps and among-site rate variations modeled with gamma distributions (shape parameter = 1). Furthermore, bootstrap resampling (n = 100) was performed to estimate statistical support for the internal branches of the NJ tree and midpoint rooting was conducted on the resulting NJ tree.

A total of 103 family-level trees were visually inspected and taxon assignments were revised in the following manner. Among U98.5 MOTUs placed within a monophyletic group consisting of a single genus, unidentified MOTUs were named after that genus, followed by "sp." with sequential numbers (e.g., *Pagrus* sp. 1, sp. 2, sp. 3...). Regarding the remaining MOTUs ambiguously placed in the family-level tree, unidentified MOTUs were named after that family, followed by "sp." with sequential numbers (e.g., Sparidae sp. 1, sp. 2, sp. 3...).

***Estimation of DNA copy numbers using quantitative PCR***

The nonlinear time series analytical tools used in the present study require a quantitative time series, namely, relative abundance data that are common for eDNA metabarcoding studies are not suitable. To estimate fish eDNA concentrations, we initially quantified the eDNA concentrations of the most common fish species in the region, Japanese black seabream (*Acanthopagrus schlegelii*), using quantitative PCR (qPCR). Metabarcoding detected the eDNA of *A. schlegelii* in 504 out of 550 water samples (91.6%), and, thus, we decided to use the sequence reads of *A. schlegelii* as the internal standard DNA of each sample.

We performed qPCR using the LightCycler® 96 System (Roche Diagnostics, Mannheim, Germany). Twenty microliters of the PCR reaction mixture comprised 2 μL of the template DNA extract, a final concentration of 900 nM each of the forward (5′–CTG TCT GCC GTC CCC TAC A–3′) and reverse (5′–TAT GGC GGC TAC GAT AAA AGG A– 3′) primers, a final concentration of 125 nM of the probe (5′–FAM-TCA GTT GAC AAC GCA ACC CTA ACC CGT AMR A–3′), and 1 × PCR master mix (FastStart Essential DNA Probes Master, Roche). The primers and probe were designed to specifically amplify a 129-bp fragment from the *A. schlegelii* mitochondrial cytochrome *b* (cyt *b*) gene (Takahashi et al., 2020). The species-specificity of the primers and probe was checked by Sasano et al. (2022). Reaction conditions were as follows: 10 min at 95˚C, 55 cycles at 95˚C for 10 s and at 60˚C for 30 s. PCR triplicates were provided for each sample. We quantified the concentrations of DNA from the calibration curve obtained from amplifying triplicates of the four standards containing $3 \times 10$, $3 \times 10^2$, $3 \times 10^3$, and $3 \times 10^4$ copies of artificial DNA fragments inserted into the target region. To check cross contamination during the PCR process, we also used triplicates of a mixture that contained 2 μL of pure water instead of template DNA. We did not detect *A. schlegelii* DNA from any FB, EB, or PCR blanks by qPCR.

We converted eDNA sequence reads obtained by eDNA metabarcoding using the quantities of *A. schlegelii* eDNA as internal standard DNAs. By dividing *A. schlegelii* sequence reads by the *A. schlegelii* eDNA quantity, we estimated how many sequence reads were generated per *A. schlegelii* eDNA copy for each sample. Note that sequence reads generated per eDNA copy may vary depending on factors such as the level of PCR inhibition and PCR amplification efficiency; however, an analog of this "internal standard DNA" method has been shown to reasonably estimate the quantity of eDNA concentrations (Ushio et al. 2018; Ushio 2022). When we did not detect any *A. schlegelii* eDNA by qPCR, we replaced the "zero" value with the minimum eDNA copy numbers of *A. schlegelii*. Similarly, when we did not detect any *A. schlegelii* eDNA sequence reads by metabarcoding, we replaced the "zero" value with the minimum eDNA sequence reads of *A. schlegelii*. These corrections enabled estimations of how many sequence reads were generated per eDNA copy for all samples.

### Supplementary Discussion

##### *Interpretations of and discussions on large information flow between fish species*

We detected statistically clear information flow between fish eDNA time series, suggesting potential interspecific interactions between fish species. Large information flow, which suggests strong interspecific interactions, may be reasonably discussed based on ecology and behavior in natural conditions. In this section, we discussed potential interspecific interactions that underlie 20 of the largest information flows between the eDNA time series described in Table S3. In the present study, information flow was typically found in species sharing a microhabitat, but not prey items. This is convincing because the eDNA times was taken at the time interval of twice a month, and thus the interactions detected by the time series analysis should also have the same time interval, which suggests that the interactions detected by our analysis could reflect behavioral interactions, rather than birth-death process.

- Effects from *Pseudoblennius marmoratus* to *Pseudolabrus eoethinus* (TE = 0.427)

  Both species live in a rocky reef habitat. *P. marmoratus* is a relatively small piscivore that preys on small pelagic fishes, while *P. eoethinus* is a large carnivore that preys on benthic crustaceans. They are likely to share a microhabitat being alarmed with common predators, i.e., large piscivores, without competition for food.

- Effects from *Pseudoblennius marmoratus* to *Gymnothorax kidako* (TE = 0.425)

  Although both are ferocious piscivores, the average size of the former is markedly smaller than that of the latter. Therefore, the former may be prey of the latter.

- Effects from *Girella punctata* to *Pseudolabrus eoethinus* (TE = 0.417)

  *G. punctata* is omnivorous and mainly feeds on algae, while *P. eoethinus* mainly feeds on epibenthos, particularly large crustaceans. The feeding activity of the former is likely to expose the prey items of the latter.

- Effects from *Pseudolabrus eoethinus* to *Gymnothorax kidako* (TE = 0.411)
- Effects from *Gymnothorax kidako* to *Pseudolabrus eoethinus* (TE = 0.36)

  These fish species are both carnivorous and may conduct joint hunting, as has been reported for the grouper *Plectropomus pessuliferus* and giant moray eel *Gymnothorax javanicus* (Bshary et al., 2006). *P. eoethinus* is a large and solitary wrasse with a high body height, making it difficult to be preyed upon by *G. kidako*.

- Effects from *Chaenogobius annularis* to *Takifugu niphobles* (TE = 0.358)

  Both species are often found in the sandy bottom substrate. The former is generally smaller than the latter. The major prey items of the former are copepods and mysids, while the latter has strong teeth to brake hard-shell benthos. The latter is highly vulnerable to ectoparasitic copepods, which may be removed by the former.

- Effects from *Girella punctata* to *Pseudoblennius marmoratus* (TE = 0.337)
- Effects from *Girella punctata* to *Gymnothorax kidako* (TE = 0.334)

  *G. punctata* feed on algae and this may provide an opportunity for *P. marmoratus* and *G. kidako* to feed on fish that hide in the algal bed.

- Effects from *Siganus fuscescens* to *Thalassoma cupido* (TE = 0.33)
- Effects from *Prionurus scalprum* to *Thalassoma cupido* (TE = 0.324)

  The feeding activities of *S. fuscescens* and *P. scalprum* on algae may enhance those of *T. cupido* on epibenthos.

- Effects from *Thalassoma cupido* to *Kyphosus bigibbus* (TE = 0.315)

  The feeding activity of *T. cupido* on epibenthos in the algal bed may facilitate *K. bigibbus* feeding on algae.

- Effects from *Siganus fuscescens* to *Kyphosus bigibbus* (TE = 0.288)

  They are both herbivores. Although they may compete for food, they may collaborate in the alert of common predators.

- Effects from *Thalassoma cupido* to *Siganus fuscescens* (TE = 0.286)

  Feeding on epibenthos by *T. cupido* may facilitate that of *S. fuscescens* on algae.

- Effects from *Chaenogobius annularis* to *Acanthopagrus schlegelii* (TE = 0.28)

  Both *C. annularis*, a small goby, and *A. schlegelii* live in shallow waters, such as the surf zone and tide pools, and, thus, these fish species may interact.

- Effects from *Chaenogobius gulosus* to *Eviota abax* (TE = 0.279)

  These fish species are goby species living in shallow waters, and, thus, they may interact.

- Effects from *Takifugu niphobles* to *Chaenogobius annularis* (TE = 0.278)

  *T. niphobles* often burrows into sand and this may provide an opportunity for *C. annularis* to feed on endobenthos, such as amphipods.

- Effects from *Thalassoma cupido* to *Prionurus scalprum* (TE = 0.269)

  The feeding of *T. cupido* on epibenthos may facilitate *P. scalprum* feeding on algae.

- Effects from *Sebastes* spp. to *Kyphosus bigibbus* (TE = 0.268)
- Effects from *Sebastes* spp. to *Siganus fuscescens* (TE = 0.266)
- Effects from *Kyphosus bigibbus* to *Sebastes* spp. (TE = 0.262)

  All of the above are combinations of the genus *Sebastes* and seaweed feeding species. *Sebastes* spp. have large eyes adapted to a low light intensity and are highly capable of detecting the approach of potential predators, while *K. bigibbus* and *S. fuscescens* feed on algae and may chase out small fish and shrimp that will become the prey of *Sebastes* spp.

It is notable that obligatory schooling species, such as Japanese Anchovy (*Engraulis japonicus*) and Silverside (*Atherion elymus*), did not have strong interactions with other species despite the high detection frequencies of their eDNA. On the other hand, strong interactions were often detected among non-schooling fish species. Therefore, this pattern suggest that interspecific interactions might complement the primary functions of fish schools, namely predator avoidance and prey detection, in these non-schooling fish species.
